# PROTAC-mediated Degradation of HIV-1 Nef Efficiently Restores Cell-surface CD4 and MHC-I Expression and Blocks HIV-1 Replication

**DOI:** 10.1101/2023.08.14.553289

**Authors:** Lori A. Emert-Sedlak, Colin M. Tice, Haibin Shi, John J. Alvarado, Sherry T. Shu, Allen B. Reitz, Thomas E. Smithgall

## Abstract

The HIV-1 Nef accessory factor is critical to the viral life cycle *in vivo* where it promotes immune escape of HIV-infected cells and viral persistence. While these features identify Nef as an attractive antiretroviral drug target, Nef lacks enzymatic activity and an active site, complicating development of occupancy-based drugs. Here we describe the development of proteolysis targeting chimeras (PROTACs) for the targeted degradation of Nef. Nef-binding compounds, based on a previously reported hydroxypyrazole core, were coupled to ligands for ubiquitin E3 ligases via flexible linkers. The resulting bivalent PROTACs induced formation of a ternary complex between Nef and the Cereblon E3 ubiquitin ligase, resulting in ubiquitylation of Nef and proteolytic degradation. Nef-directed PROTACs efficiently rescued Nef-mediated MHC-I and CD4 downregulation in T cells and suppressed HIV-1 replication in donor PBMCs. Targeted degradation of Nef is anticipated to reverse all HIV-1 Nef functions and may help restore adaptive immune responses against HIV-1 reservoir cells *in vivo*.

## INTRODUCTION

The *nef* genes of the primate lentiviruses HIV-1, HIV-2 and SIV encode unique membrane-associated proteins with key roles in viral replication, persistence, and AIDS progression. HIV-1 Nef is expressed at high levels soon after infection and interacts with diverse host cell proteins to enhance viral infectivity and promote immune escape. Some of the best-studied Nef partner proteins include the endocytic adaptor proteins AP-1 and AP-2 which are required for downregulation of MHC-I, CD4 and other viral receptors^1,2^ as well as the SERINC5 restriction factor.^3-5^ Nef also binds and activates non-receptor protein-tyrosine kinases^6^ and regulators of the actin cytoskeleton^7^ to promote viral transcription and release.

Non-human primate studies provided some of the first evidence that Nef is essential for viral pathogenesis and the development of AIDS. Nef-defective SIV replicates poorly in rhesus macaques, resulting in low viral loads and delayed disease onset.^8^ These studies mirror reports of individuals infected with Nef-defective HIV-1 in which viral loads remain low in the absence of antiretroviral therapy.^9-11^ Similar observations have been made in HIV-1 infected humanized immune system mice, in which immunodeficient animals are reconstituted with human CD4^+^ T cells and other host cell targets for HIV-1.^12,13^ In these animals, wild-type HIV-1 infection results in plasma viremia and depletion of CD4^+^ T cells while Nef-defective HIV-1 replicates poorly and does not cause CD4^+^ T cell loss.^11,14^ These animal model and HIV patient data establish a central role for Nef in HIV-1 pathogenesis and support the development of Nef antagonists as new therapeutic approach against HIV/AIDS.^15,16^

Several studies have reported drug discovery efforts targeting HIV-1 Nef.^15,16^ One approach involved high-throughput screening of chemical libraries for inhibitors of Nef-dependent activation of a protein kinase partner, the Src-family kinase, Hck.^17,18^ This screen identified a unique Nef-binding compound based on a hydroxypyrazole core that inhibited Nef-dependent enhancement of HIV-1 infectivity, replication, and partially reversed Nef-mediated MHC-I downregulation.^17,19^ Nef is well-known to inhibit cell-surface display of MHC-I in complex with HIV-1 antigenic peptides thereby preventing detection by cytotoxic T lymphocytes.^1^ In this way, Nef interferes with clearance of the virus from the infected host, and may contribute to establishment and maintenance of the persistent viral reservoir.^20,21^ The original hydroxypyrazole Nef inhibitor identified by high-throughput screening as well as first-generation analogs^22^ based on this structure restored MHC-I to the surface of latently HIV-infected CD4^+^ T cells *in vitro*.^19^ Moreover, when inhibitor-treated cells were co-cultured with autologous CD8^+^ T cells expanded in the presence of HIV-1 antigenic peptides, the CD8^+^ T cells became activated and displayed CTL responses against the infected CD4^+^ target cells.^19^ This proof-of-concept study strongly suggests that Nef inhibitors may enhance CTL-mediated responses *in vivo* to help clear the latent viral reservoir.

In more recent work, our team synthesized an extensive series of hydroxypyrazoles that bind tightly to Nef by surface plasmon resonance (SPR) and are active against multiple Nef functions *in vitro*.^23^ However, Nef lacks an active site, complicating medicinal chemistry optimization due to the lack of correlation between Nef binding affinity *in vitro* and antiretroviral activity in cell-based systems. In addition, these occupancy-based Nef inhibitors show relatively modest efficacy in terms of cell-surface MHC-I rescue.^23^ This observation may reflect a stoichiometric inhibitor requirement to prevent Nef association with the C-terminal tail of MHC-I and the AP-1 endocytic adaptor protein responsible for its downregulation.^24^ In the present study, we addressed these limitations by repurposing existing Nef-binding compounds as bivalent <u>Pro</u>teolysis <u>Ta</u>rgeting <u>C</u>himeras (PROTACs) for the targeted degradation of Nef as illustrated in Figure 1. A major advantage of the PROTAC approach is that it requires only a selective binder of the target protein (Nef in this case) and not a functional inhibitor *per se*.^25,26^ We synthesized a library of Nef-directed PROTAC candidates that join existing hydroxypyrazole Nef-binding compounds to ligands for the Cereblon (CRBN) and Von Hippel-Lindau (VHL) components of ubiquitin E3 ligases. Using orthogonal cell-based assays for Nef ubiquitylation, degradation, and function, we identified promising Nef-directed PROTACs which target the CRBN E3 ligase pathway. These compounds induce Nef degradation and efficiently reverse Nef-mediated MHC-I and CD4 downregulation. Nef PROTACs also demonstrate potent antiretroviral activity in HIV-infected primary cells, supporting further development for testing in the context of HIV-1 reservoir reduction *in vivo*. More broadly, these results encourage the development of PROTACs against viral proteins which are intrinsically difficult to target with traditional occupancy-based approaches.^27,28^

**Figure 1.**
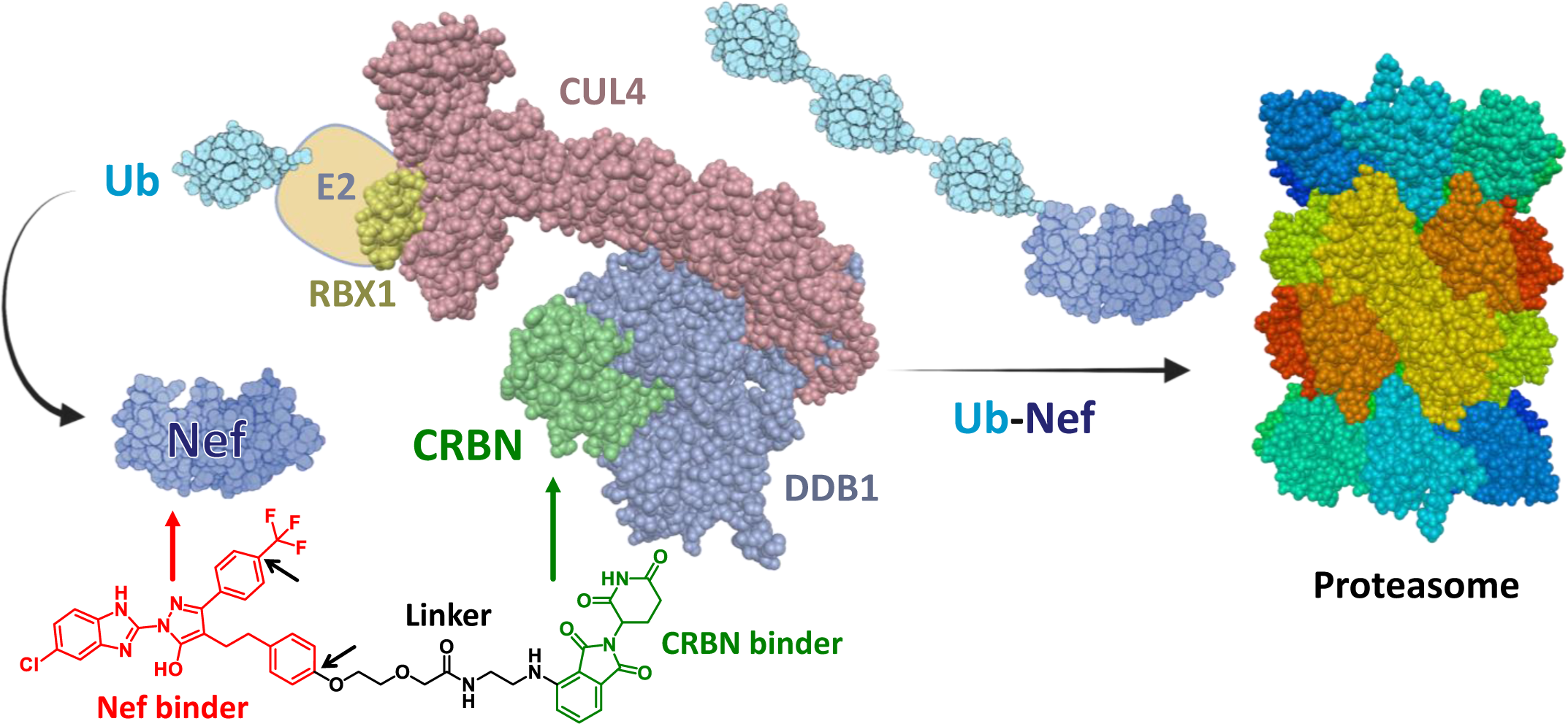
Targeted degradation of HIV-1 Nef by a CRBN-based PROTAC. The Cereblon (CRBN) ubiquitin E3 ligase complex (*left*) is a large multiprotein structure composed of RING-box protein 1 (RBX1), Cullin4 (CUL4), DNA damage binding protein 1 (DDB1), CRBN and an E2 subunit conjugated to ubiquitin (Ub). Heterobifunctional Nef PROTACs promote formation of a ternary complex between the HIV-1 Nef protein using existing hydroxypyrazole Nef-binding compounds (*red*) and the CRBN E3 complex via a CRBN ligand (exemplified by thalidomide, green). Ternary complex formation induces polyubiquitination of Nef and subsequent proteasomal degradation. The Nef PROTAC shown is analog **FC-13182**; favored positions for linker attachment on the Nef-binding moiety are indicated by the *short black arrows*.

## RESULTS

### Synthesis of PROTACs targeting HIV-1 Nef

To create Nef-directed PROTACs, we first attached prototypical PEG linkers to various positions on previously developed Nef-binding compounds.^23^ Screening of these probe compounds for Nef binding by SPR indicated that the *para* positions of the two phenyl rings tolerate linker attachment without substantial loss in binding affinity (arrows in Figure 1). We then coupled ligands for ubiquitin E3 ligase components (CRBN or VHL) with linkers of varying lengths and chemistries attached to these two *para* positions. A total of 102 unique Nef-directed PROTACs were synthesized, with 77 analogs linked to CRBN ligands (thalidomide and lenalidomide) and 25 linked to VHL ligands. Details of the synthetic procedures and analytical characterization of active analogs are presented in the Supplemental Information.

### Assessment of PROTAC induction of Nef ubiquitylation by NanoBRET Assay

Candidate Nef-directed PROTACs were first screened for induction of Nef ubiquitylation in a cell-based NanoBRET assay (Promega; Figure 2A). This assay consists of two key components: 1) HIV-1 Nef fused via its C-terminus to the small luciferase protein, nano-Luc (Nef-nLuc), and 2) Ubiquitin fused to the self-labeling fluorescent protein known as the Halo-Tag (Ub-Halo).^29^ The Halo-Tag is a modified bacterial haloalkane dehalogenase that forms a covalent adduct in the presence of a chloroalkane coupled to a fluorogenic acceptor (NanoBRET 618 in this case). Nef-nLuc and Ub-Halo were co-expressed in 293T cells for 18 hours, followed by addition of each PROTAC analog plus the Ub-Halo ligand. Twenty-four hours later, the nano-Luc substrate was added, and the donor (nLuc; 460 nm) and acceptor (HaloTag; 618 nm) signals were recorded. Proximity of Nef-nLuc and Ub-Halo resulting from successful ubiquitylation of Nef results in a BRET signal. Each PROTAC was assayed in quadruplicate, and the resulting data were corrected for background and expressed as 618 nm to 460 nm fluorescence ratios (BRET signal for Ub incorporation normalized to Nef-nLuc levels). BRET ratios were then normalized to the DMSO control wells and used to calculate z-scores based on the normalized ratios for all analogs tested (Figure 2B). All PROTACs were also assayed in quadruplicate in control assays with the Halo-Tag alone. Control assays produced BRET ratios significantly lower than those observed in the presence of Ub-Halo, demonstrating the dependence of the readout on Nef ubiquitylation. Overall, this analysis identified eleven PROTACs that increased Nef ubiquitylation by at least 1.5 standard deviations above the mean (z-score > 1.5). These analogs were advanced to orthogonal assays for induction of Nef degradation and inhibition of Nef function. Analog **FC-13887** (z-score = 1.3) was also carried forward due to its activity in the receptor rescue assay described below. Eleven of the 12 active analogs targeted CRBN in this assay, while only a single VHL-based PROTAC showed significant activity (**FC-14373**). NanoBRET assay data for the twelve active Nef PROTAC analogs, including the negative controls, are summarized in the Supplemental Information, Figure S1.

**Figure 2.**
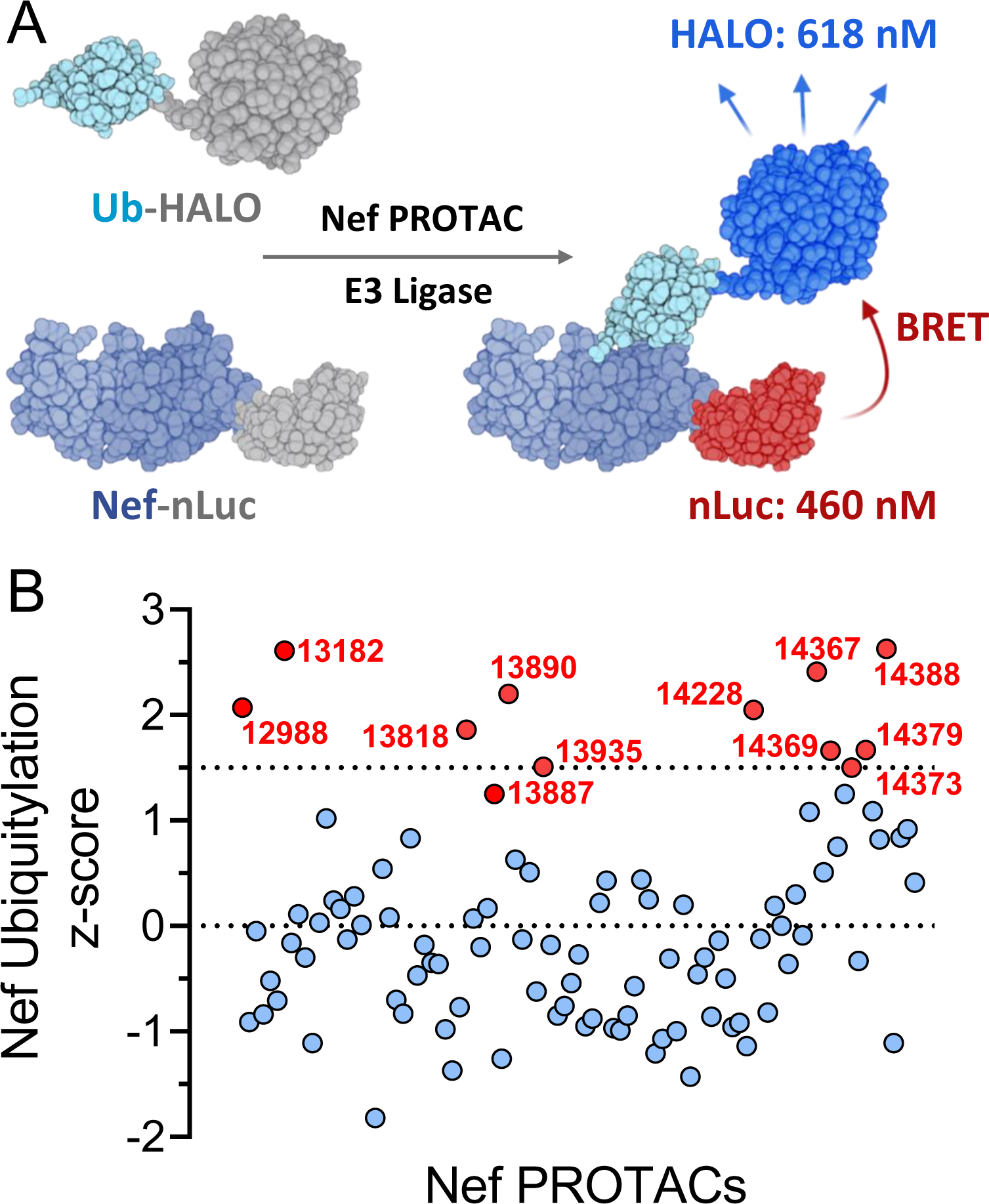
NanoBRET assay for PROTAC-mediated ubiquitination of HIV-1 Nef. A) Assay principle. Nef is fused to nano-Luciferase (Nef-nLuc) and co-expressed with a ubiquitin-Halo tag fusion protein (Ub-Halo) in 293T cells. PROTACs promote ligation of Ub-Halo to Nef-nLuc, which is detected by bioluminescence resonance energy transfer (BRET) to the Halo Tag. B) Assessment of candidate Nef PROTACs in the NanoBRET assay. Each compound was assayed in quadruplicate and the average 618 nm to 460 nm fluorescence ratios (BRET signal for Ub incorporation normalized to Nef-nLuc levels) were normalized to the DMSO control and are presented as z-scores ± SD (error bars smaller than data points). PROTACs with z-scores ≥ 1.5 (numbered red points) along with analog **FC-13887** were advanced to orthogonal assays for Nef degradation and inhibition of Nef function. z-score = (x - µ)/σ, where x = each individual value, µ = mean value, and σ = the standard deviation.

### T cell-based assay for PROTAC-induced Nef degradation and cell surface receptor downregulation

For these experiments, the T cell line CEM-T4 was stably transduced with a doxycycline (Dox)-inducible expression vector for a Nef-eGFP (enhanced GFP) fusion protein. In the presence of Dox, Nef-eGFP is expressed leading to downregulation of CD4 and MHC-I from the cell surface which are quantified by flow cytometry (Figure 3A). CEM/Nef-eGFP cells were incubated with each of the active PROTACs from the Nef-Ub NanoBRET screen at a final concentration of 3 µM or DMSO as control, and Nef expression was induced by the addition of Dox. After 24 h, treated cells were analyzed for reversal of cell-surface CD4 and MHC-I downregulation. Six of the 12 Nef PROTACs tested restored cell-surface CD4 expression by more than 50% while seven PROTACs restored cell-surface MHC-I by 50% or more as well (Figure 3B). These observations represent a substantial improvement over the previous occupancy-based Nef inhibitors on which the PROTACs are based, which reversed Nef-dependent MHC-I downregulation in the 5-15% range under similar conditions.^23^

**Figure 3.**
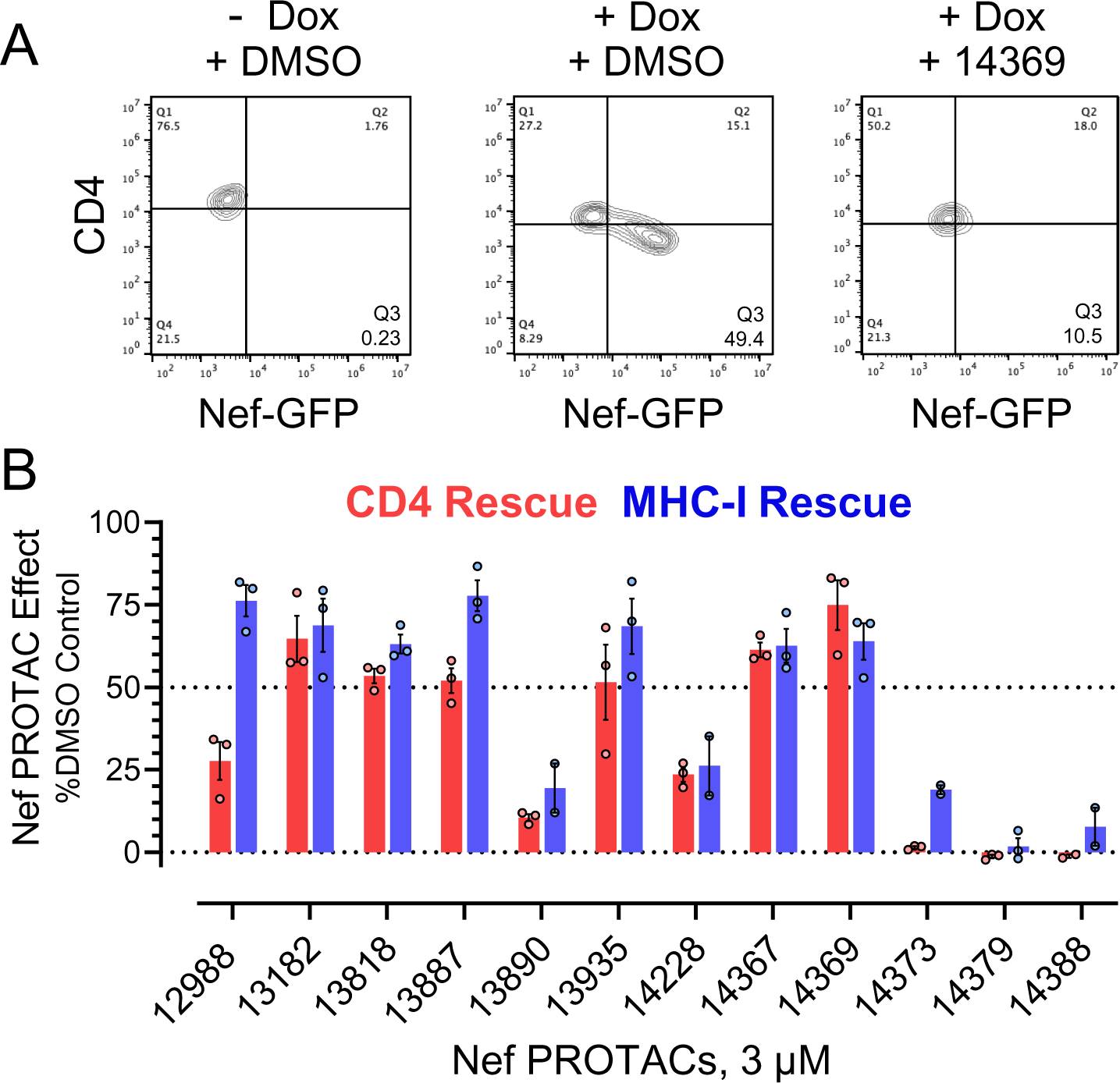
Nef PROTAC treatment restores cell surface CD4 and MHC-I expression in T cells. The human T cell line CEM-T4 was engineered to express a Nef-eGFP fusion protein under the control of a doxycycline (Dox) inducible promoter. In the absence of Dox, these cells express endogenous CD4 and MHC-I on their surface; addition of Dox induces Nef-eGFP expression which leads to receptor downregulation. A) Representative flow cytometry result with Nef PROTAC **FC-14369** and cell surface CD4 staining. B) Active Nef PROTACs from the NanoBRET ubiquitination assay were screened for cell surface receptor rescue in triplicate. Bar height indicates the mean value ± SE; individual data points are also shown. The structures of the analogs with little to no activity in this assay (**FC-13890**, **FC-14373**, **FC-14379**, **FC-14388**) are shown in the Supplemental Information, Figure S2.

To determine whether PROTAC-mediated rescue of cell-surface receptor expression was due to degradation of the Nef protein, we also calculated levels of Nef-eGFP in each treated cell population by flow cytometry. PROTACs that restored cell surface receptor expression also induced significant loss of Nef-eGFP in the cell population, with values for percent loss ranging from 15% to more than 50% (Figure 4A). Rescue of cell surface CD4 expression showed a significant linear correlation with Nef-eGFP loss, with the degree of cell surface receptor rescue induced by each PROTAC tracking with the extent of Nef-eGFP reduction (Figure 4B). On the other hand, rescue of cell-surface MHC-I expression showed a non-linear correlation with Nef-eGFP loss, with the activity of the most active compounds plateauing at around 75% of control. As an independent measure of PROTAC-mediated loss of Nef expression, we also performed quantitative immunoblot analysis of PROTAC-treated CEM cell lysates using antibodies specific for Nef as well as Actin as a normalizing control (Figure 4C). This experiment confirmed PROTAC-dependent loss of Nef protein expression, with values decreasing from 65% to more than 95% relative to the DMSO-treated control (Figure 4D). These results provide independent evidence that the PROTACs induce Nef degradation, thereby restoring cell-surface expression of receptors essential for immune system recognition of HIV-infected cells.

**Figure 4.**
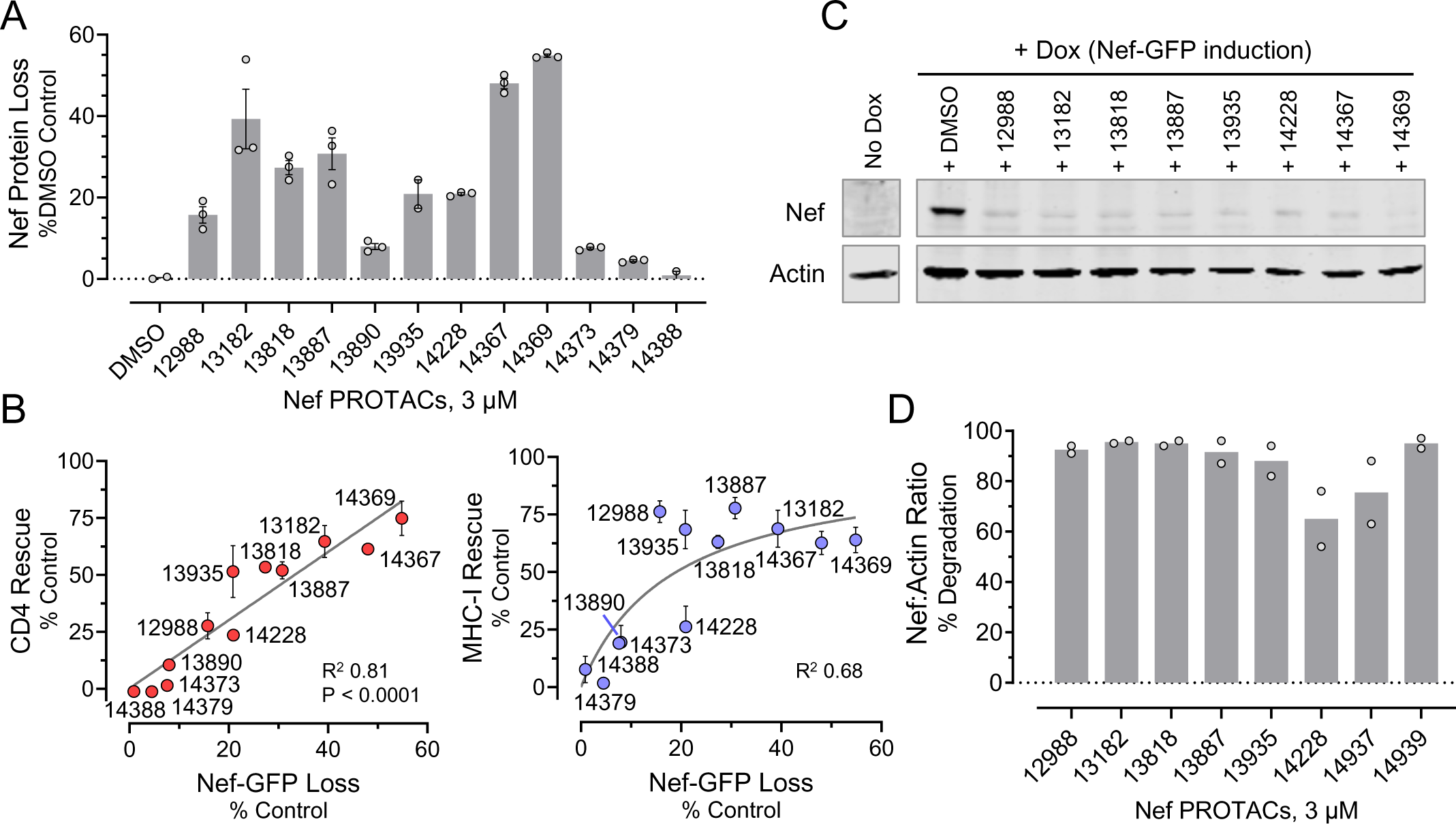
Assessment of PROTAC-mediated Nef protein loss. A) Flow cytometry of Nef-eGFP protein loss. CEM/Nef-eGFP cells were treated with doxycycline to induce expression of Nef-eGFP under conditions that result in a moderate level of positive cells by flow cytometry (see Figure 3A). Triplicate cultures of cells were treated with the Nef PROTAC analogs indicated at a final concentration of 3 µM, and 24 h later the percent of cells showing loss of Nef-eGFP protein expression were calculated relative to the DMSO controls and are presented as the mean value ± SE; individual data points are also shown. B) Correlation analysis of cell-surface CD4 rescue vs. Nef-eGFP protein loss (red data points, left) and MHC-I rescue vs. Nef-eGFP protein loss (blue data points, right). CD4 rescue was best-fit by linear regression, while MHC-I rescue showed a plateau effect. C) Immunoblot analysis. Cells expressing Nef-eGFP were treated as in part A with the eight active PROTACs, and lysates were prepared 48 h later for immunoblot analysis with Nef and Actin antibodies. A representative blot is shown. D) Immunoblot analysis was performed in duplicate, and band intensities were quantified by LI-COR infrared imaging and used to calculate Nef to Actin protein expression ratios. The bar graph shows the mean value for each ratio along with the individual values. The structures of the analogs with little to no activity in this assay (**FC-13890**, **FC-14373**, **FC-14379**, **FC-14388**) are shown in the Supplemental Information, Figure S2.

Structures of the eight most active Nef PROTACs are presented in Figure 5. Six analogs use thalidomide as the CRBN-targeting moiety while the other two are based on lenalidomide. In five analogs, the CRBN-recruiter is attached to the phenyl group of the Nef ligand analog via the *para* position (position ‘**x**’) while three used the *para* position of the phenylethyl group (position ‘**y**’). A wide range of linker lengths was tolerated, ranging from just three atoms (**FC-14367**) to more than 20 (**FC-14228**). These observations suggest multiple starting points for the synthesis of analogs with optimized physicochemical and pharmacological properties for further development. The structures of the analogs with little to no activity in the receptor downregulation and degradation assays (**FC-13890**, **FC-14373**, **FC-14379**, **FC-14388**) are shown in the Supplemental Information, Figure S2.

**Figure 5.**
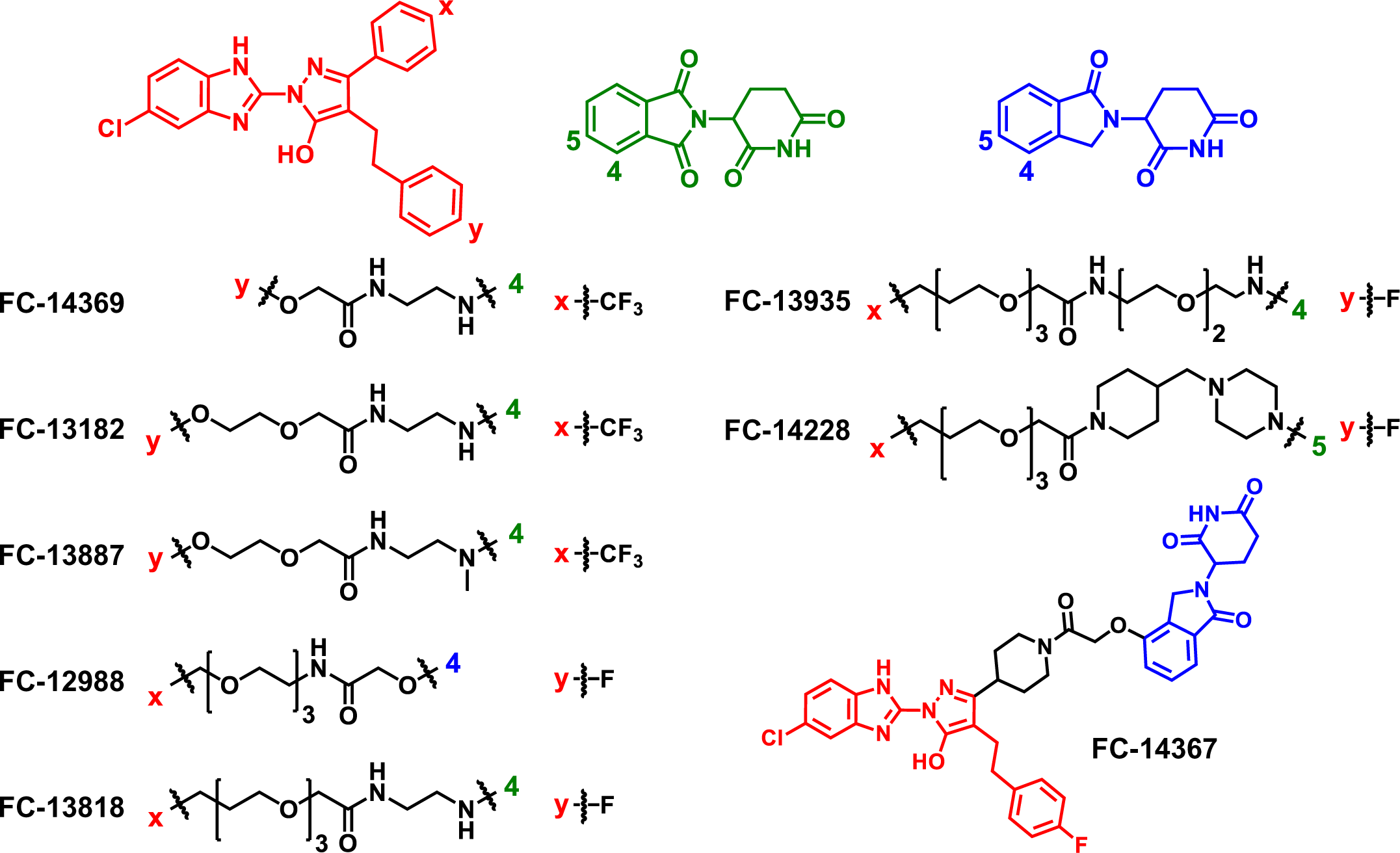
Structures of active Nef PROTACs. The Nef-targeting moiety (red, upper left) provides two positions on the phenyl rings for linker attachment (**x** and **y**). CRBN ligands included analogs of thalidomide (green, center) and lenalidomide (blue, right) that were attached to the isoindoline ring at positions 4 and 5. Linker compositions are shown below adjacent to each analog number. The complete structure of **FC-14367** is also provided.

### Assessment of direct PROTAC binding to Nef and CRBN by SPR

To confirm interaction of each bivalent PROTAC analog with both Nef and CRBN, we used our established SPR assay for small molecule-protein interactions.^23^ Eight recombinant Nef proteins representative of multiple M-group HIV-1 variants, SIV Nef mac239, and the thalidomide-binding domain of CRBN were expressed in bacteria and purified to homogeneity. The recombinant proteins were immobilized on a carboxymethyl dextran hydrogel biosensor chip, and solubilized PROTACs were injected over a range of concentrations in triplicate. Following a dissociation phase, the resulting sensorgrams were fitted with a 1:1 Langmuir binding model and K_D_ values were calculated from the rate constants and the relationship, K_D_ = k_d_/k_a_. Representative SPR sensorgrams for active Nef PROTACs **FC-12988** and **FC-13182** are shown in Figure 6 to illustrate differences in the binding kinetics. While both PROTACs bound to the CRBN thalidomide-binding domain to a similar extent, **FC-12988** associated more slowly than **FC-13182** and showed a comparatively slow dissociation phase. The reverse was true with HIV-1 Nef (NL4-3 variant), in which **FC-12988** showed more rapid association and release compared to **FC-13182**. These kinetic differences may in part reflect the alternative points of linker attachment on the Nef-binding moiety. The shape of the PROTAC sensorgrams with HIV-1 Nef closely resemble those reported previously for structurally related Nef-binding components alone.^23^ However, the extent of binding (RU value) was higher at each PROTAC concentration, which likely reflects the higher formula weight of the PROTACs. Regardless, both compounds effectively induced Nef degradation and restored CD4 and MHC-I expression to the surface of T cells.

**Figure 6.**
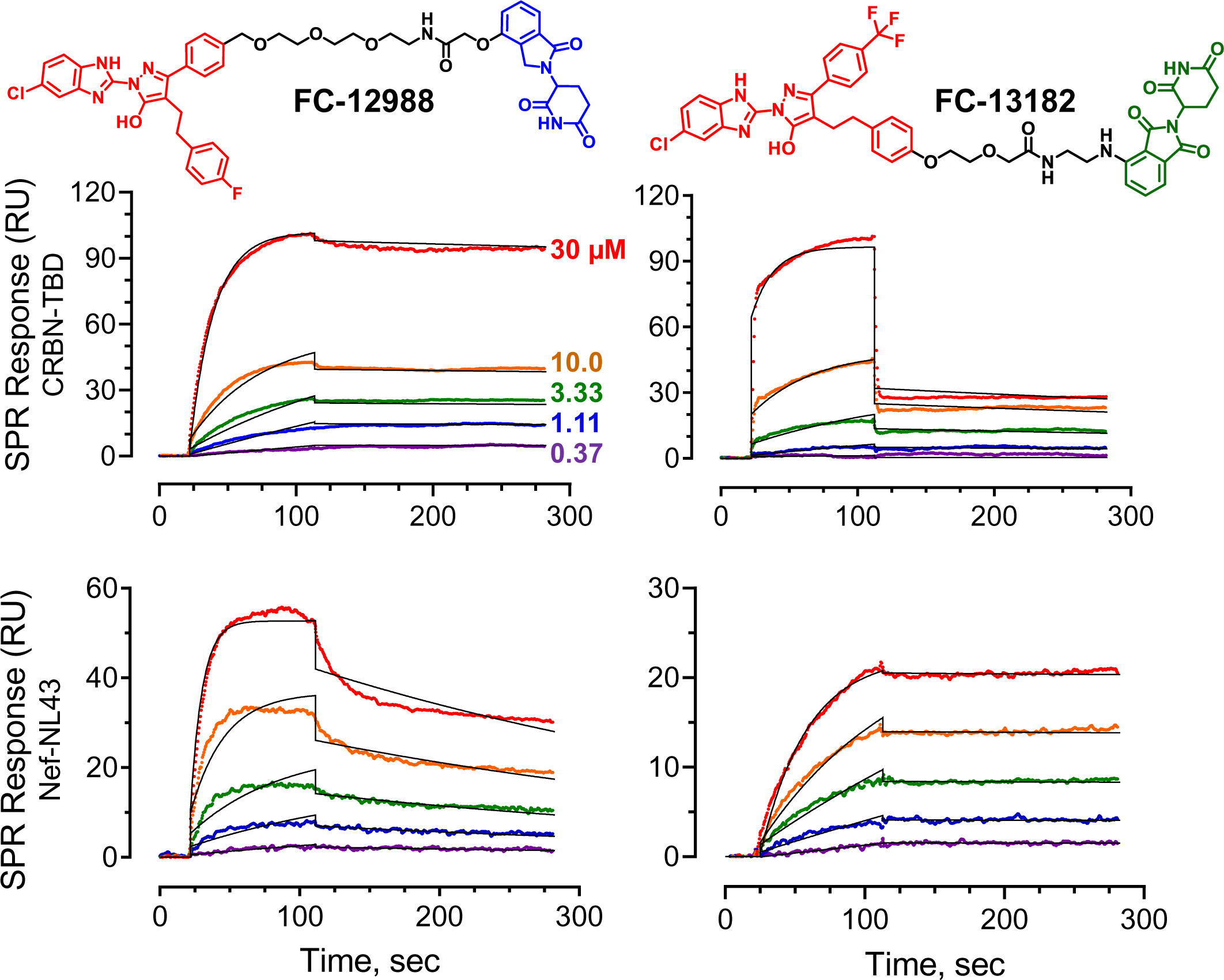
Representative SPR sensorgrams for Nef PROTACs. The thalidomide-binding domain of Cereblon (CRBN-TBD) and full-length Nef (NL4-3 variant) were expressed in *E. coli* and purified to homogeneity. Each protein was immobilized on one channel of a carboxymethyl dextran biosensor chip, and the two Nef PROTAC analogs **FC-12988** and **FC-13182** (structures at top; Nef-binding moiety in red, CRBN-binding ligands thalidomide and lenalidomide are shown in green and blue, respectively) were injected over the range of concentrations shown in the upper left sensorgram. Protein-ligand interaction was followed for 90 s, followed by a 180 s dissociation phase. The resulting data were fit to a 1:1 Langmuir binding model, and K_D_ values were calculated from the resulting association and dissociation rate constants (K_D_ values are summarized in Table 1). Each concentration was tested in triplicate, and individual traces are shown with the data shown in color and the fitted curves superimposed in black. RU, SPR response units.

**Table 1.** Direct binding of active Nef PROTACs to multiple HIV-1 Nef variants, SIV Nef, and the thalidomide binding domain of Cereblon (CRBN) by SPR. Nef proteins include the well-characterized B-subtype variants SF2 and NL4-3 and the C-clade transmitted/founder (T/F) variant, C/z3618m, as well as SIV Nef mac239. K_D_ values (M) were calculated from the association and dissociation rate constants using the relationship, K_D_ = k*_d_*/k*_a_*.

| Analog<br>FC- | Representative M-group HIV-1 Nef Subtypes |  |  |  |  |  |  |  | SIV Nef | CRBN |
| --- | --- | --- | --- | --- | --- | --- | --- | --- | --- | --- |
|  | B (NL4-3) | B (SF2) | A1 | C (T/F) | F1 | G | J | K |  |  |
| 12988 | $6.06 \times 10^{-7}$ | $6.93 \times 10^{-7}$ | $1.04 \times 10^{-6}$ | $2.30 \times 10^{-5}$ | $6.77 \times 10^{-5}$ | $4.05 \times 10^{-7}$ | $6.29 \times 10^{-8}$ | $4.50 \times 10^{-7}$ | $6.88 \times 10^{-7}$ | $5.14 \times 10^{-7}$ |
| 13182 | $5.42 \times 10^{-8}$ | $5.55 \times 10^{-7}$ | $9.69 \times 10^{-7}$ | $3.51 \times 10^{-7}$ | $1.96 \times 10^{-5}$ | $1.48 \times 10^{-7}$ | $3.99 \times 10^{-8}$ | $1.65 \times 10^{-7}$ | $2.27 \times 10^{-8}$ | $1.35 \times 10^{-8}$ |
| 13818 | $5.39 \times 10^{-7}$ | $1.36 \times 10^{-6}$ | $1.83 \times 10^{-6}$ | $6.44 \times 10^{-6}$ | $6.02 \times 10^{-5}$ | $6.65 \times 10^{-7}$ | $1.55 \times 10^{-8}$ | $1.48 \times 10^{-6}$ | $1.78 \times 10^{-7}$ | $4.91 \times 10^{-8}$ |
| 13887 | $1.56 \times 10^{-7}$ | $6.12 \times 10^{-7}$ | $9.24 \times 10^{-7}$ | $4.75 \times 10^{-7}$ | $6.91 \times 10^{-5}$ | $4.90 \times 10^{-7}$ | $4.43 \times 10^{-7}$ | $4.75 \times 10^{-7}$ | $2.04 \times 10^{-8}$ | $1.83 \times 10^{-8}$ |
| 13935 | $3.09 \times 10^{-8}$ | $1.26 \times 10^{-6}$ | $1.39 \times 10^{-6}$ | $8.58 \times 10^{-7}$ | $3.99 \times 10^{-6}$ | $4.44 \times 10^{-7}$ | $3.52 \times 10^{-7}$ | $1.97 \times 10^{-7}$ | $1.81 \times 10^{-8}$ | $3.86 \times 10^{-9}$ |
| 14228 | $9.66 \times 10^{-7}$ | $6.94 \times 10^{-7}$ | $7.49 \times 10^{-7}$ | $2.63 \times 10^{-6}$ | $1.44 \times 10^{-6}$ | $1.60 \times 10^{-6}$ | $6.10 \times 10^{-7}$ | $8.11 \times 10^{-7}$ | $1.70 \times 10^{-8}$ | $2.36 \times 10^{-8}$ |
| 14367 | $5.25 \times 10^{-7}$ | $9.32 \times 10^{-7}$ | $9.31 \times 10^{-7}$ | $8.08 \times 10^{-7}$ | $9.35 \times 10^{-7}$ | $6.38 \times 10^{-7}$ | $4.57 \times 10^{-7}$ | $4.16 \times 10^{-7}$ | $4.48 \times 10^{-7}$ | $4.25 \times 10^{-7}$ |
| 14369 | $5.76 \times 10^{-9}$ | $5.32 \times 10^{-7}$ | $5.78 \times 10^{-7}$ | $6.59 \times 10^{-8}$ | $1.83 \times 10^{-7}$ | $1.50 \times 10^{-7}$ | $2.53 \times 10^{-8}$ | $4.35 \times 10^{-8}$ | $5.14 \times 10^{-9}$ | $1.14 \times 10^{-9}$ |

The eight most active PROTACs from the CEM T cell assay bound to all Nef variants tested as well as the CBRN thalidomide-binding domain with nM to low µM affinity in almost every case (Table 1), suggesting that PROTACs based on the hydroxypyrazole Nef-targeting moiety are widely active against M-group HIV-1 variants responsible for the pandemic. All eight PROTACs also interacted with SIV Nef_mac239_ in the nanomolar range, suggesting that these analogs may enable exploration of the effects of targeted Nef degradation in SIV/Rhesus macaque models of HIV-1 latency (see Discussion). Control SPR data show that the thalidomide analogs used for PROTAC synthesis bind to HIV-1 and SIV Nef proteins with low potency (10-70 µM range) relative to the CRBN thalidomide binding domain (1-8 µM range; Table S1).

### Active Nef PROTACs stabilize ternary Nef-CRBN complexes in vitro

Targeted protein degradation requires PROTAC-mediated formation of a ternary complex between the protein target, the bivalent PROTAC ligand, and the E3 ligase. To model this mechanism *in vitro*, we incubated recombinant Nef (NL4-3 variant) and the CRBN thalidomide-binding domain in the presence and absence of the active Nef PROTAC analog, **FC-13818**. In the absence of the PROTAC, Nef and CRBN coeluted from a size-exclusion column (Figure 7A); note that the retention volumes of the individual proteins are the same, so the mixture elutes as a single peak (Figure S3). When preincubated in the presence of the PROTAC, however, a new peak of smaller retention volume (higher molecular weight) was observed, and the height of this peak increased in proportion to the concentration of the PROTAC added to the mixture (Figure 7B). Similar results were observed with PROTACs **FC-12988** and **FC-13887** (Figure S3). These results provide evidence that active Nef PROTACs induce ternary complexes of Nef with CRBN and are consistent with the results from the NanoBRET and CEM-T4 assays.

**Figure 7.**
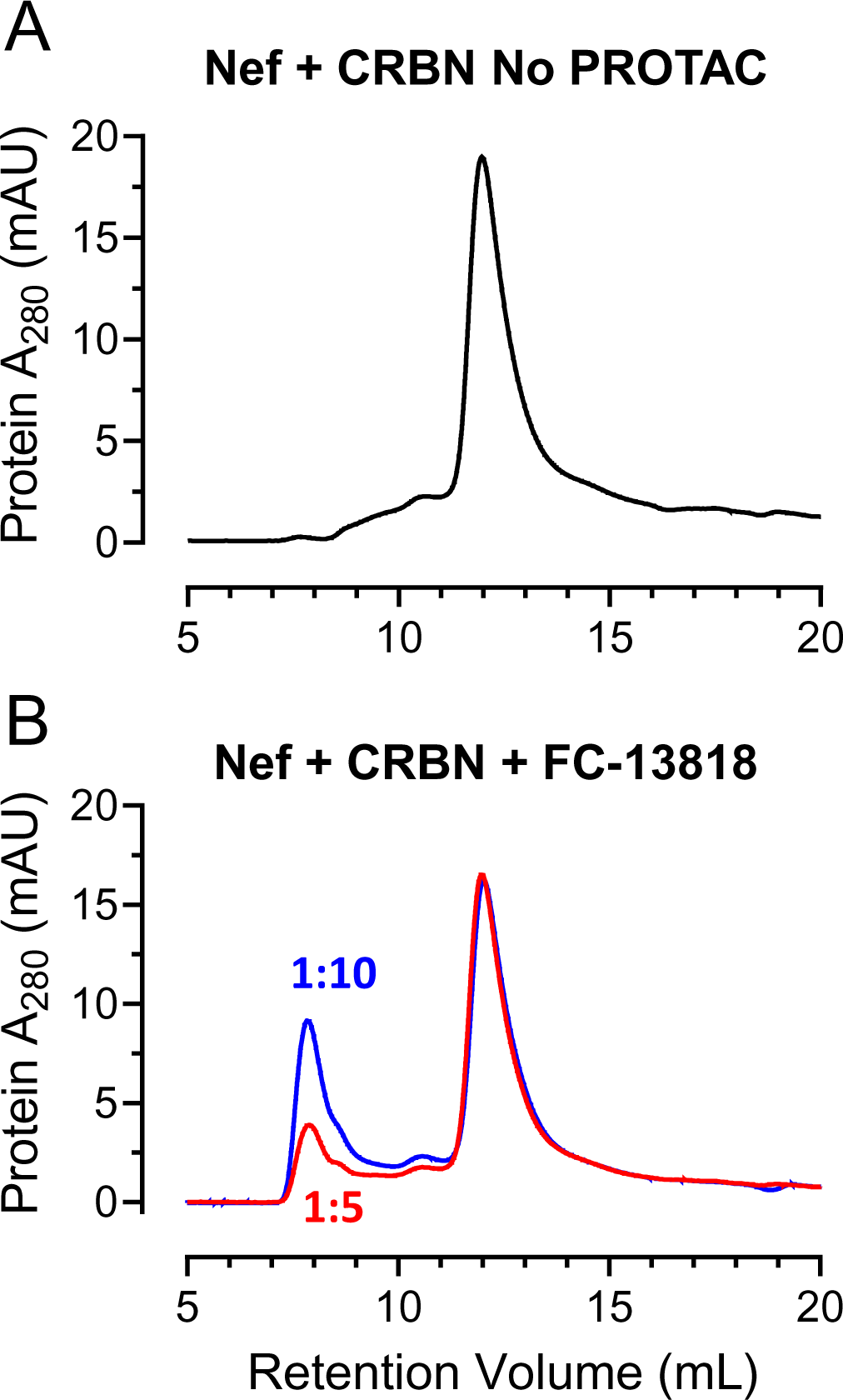
Active PROTAC FC-13818 induces Nef-CRBN protein complex *in vitro*. A) Recombinant purified Nef and the CRBN ligand-binding domain were mixed in equimolar proportions and analyzed by size exclusion chromatography (Superdex 75). The mixture elutes as a single peak as the individual proteins have similar retention volumes. B) The protein mixture from part A was incubated with Nef PROTAC **FC-13818** in the molar ratios shown at 4 °C for 20 min prior to SEC. Results with additional PROTACs and controls are shown in Figure S3.

### PROTACs block Nef-mediated enhancement of HIV-1 replication in donor PBMCs

HIV-1 Nef is well known to enhance viral replication in donor PBMCs *in vitro* and is essential for high viral loads *in vivo* (see Introduction). Our previous studies have shown that the hydroxypyrazole Nef-binding compounds used to create the PROTACs possess significant antiretroviral activity.^23^ To determine whether antiretroviral activity is retained by the Nef PROTACs, we tested the most active analogs in HIV-infected PBMCs from normal donors. PBMCs were infected with wild-type and Nef-defective HIV-1*_NL4-3_* under conditions where the presence of Nef enhanced replication by 3.3-fold consistent with prior work (Figure 8A).^23^ Cultures infected with wild-type HIV-1 were incubated with each PROTAC at a final concentration of 1 µM, and the extent of replication was assessed using a p24 Gag AlphaLISA assay 48 h later. All eight PROTACs showed antiviral activity, with six of eight analogs completely suppressing Nef-dependent enhancement of viral replication (Figure 8A). To assess cytotoxicity, uninfected PBMCs were incubated with each Nef PROTAC at 1 µM for 48 h, followed by assessment of viability with the CellTiter-Blue Assay. PBMCs treated with seven of the eight PROTACs showed viability of 90% or more compared to the DMSO control (Figure 8B). The one exception was **FC-13818**, which reduced viability by 20%; this observation may partially explain why this analog reduced HIV-1 replication below the level of the ΔNef mutant.

**Figure 8.**
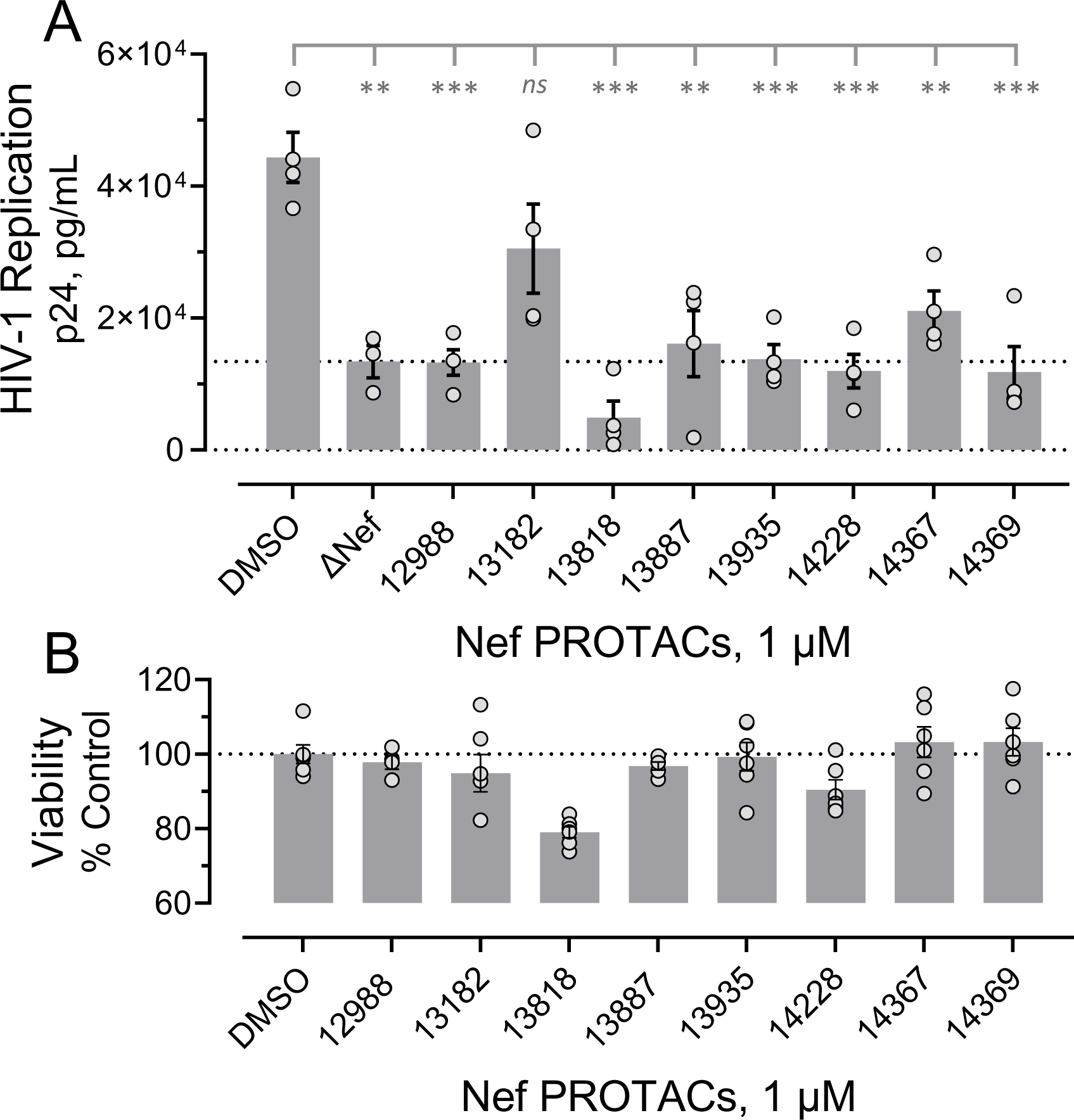
PROTACs inhibit Nef-dependent enhancement of HIV-1 replication in primary cells. A) Donor PBMCs were infected with wild-type HIV-1*_NL4-3_* (DMSO control), a Nef-defective mutant (ΔNef), or wild-type virus in the presence of the Nef PROTACs shown at a final concentration of 1 µM. Input virus was 2,500 pg HIV p24 Gag per well. Replication was assayed by p24 Gag AlphaLISA 4 days later. Six independent determinations were assayed for each condition, and the highest and lowest p24 values were removed. Each bar indicates the mean ± SE of the remaining values with the individual data points shown. The dotted line indicates the mean value for the ΔNef control. Statistical significance was determined by Student’s t test between the DMSO control and ΔNef as well as each PROTAC treatment group; **, p < 0.01; ***, p < 0.001; *ns*, not significant. B) Viability of uninfected PBMCs was determined with each PROTAC at 1 µM after 4 days using the Cell Titer Blue assay (n = 6 wells/condition). The dotted line indicates 100% viability based on the DMSO control.

## DISCUSSION

In this report, we present the synthesis and characterization of CRBN-based PROTACs that induce HIV-1 Nef proteolytic degradation via ubiquitination. Targeted degradation of Nef resulted in robust recovery of cell-surface MHC-I and CD4 expression in a T cell system while potently suppressing Nef-mediated enhancement of HIV-1 replication in primary cells. Nef is well known to interact with diverse host cell proteins to enhance the viral life cycle, promote immune escape, and support viral persistence.^1,6,15,16,21^ Nef lacks an active site and uses multiple structural motifs to recruit cellular signaling partners, complicating traditional occupancy-based antiretroviral drug development. The PROTAC approach detailed here may circumvent this issue, as targeted degradation is likely to antagonize all Nef functions while limiting side effects from non-specific activities.

The active Nef PROTACs share a Nef-targeting moiety based on a hydroxypyrazole core joined to a benzimidazole substituent (Figure 5). Previous studies of these Nef antagonists showed low nanomolar inhibition of HIV-1 replication in primary cells,^23^ an effect likely attributed to interference with Nef-activated kinases linked to viral transcription.^6,15^ Our findings that these same Nef antagonists, in the context of the PROTACs described here, cause ubiquitylation and degradation of Nef provides strong evidence for their direct interaction with Nef in a cellular context. This observation is also consistent with the ability of these PROTAC analogs to bind directly to recombinant Nef by SPR and to induce formation of ternary complexes with the CRBN thalidomide-binding domain *in vitro* (Figures 6 and 7). Importantly, Nef PROTACs based on the hydroxypyrazole targeting moiety bound to a diverse panel of HIV-1 Nef variants as well as SIV Nef (Table 1), suggesting that they will be broadly active against multiple HIV-1 strains through a shared structure feature within Nef. Alignment of the folded cores of HIV-1 and SIV Nef proteins previously determined by X-ray crystallography reveals strong structural homology in this region, along with multiple exposed lysine residues for ubiquitin attachment. This observation suggests that some of the PROTACs described here may be suitable for testing in non-human primate models of SIV latency. Figure S4 provides amino acid and structural alignments of representative HIV-1 and SIV Nef proteins (NL4-3 and mac239, respectively) and highlights the locations of surface lysine residues that represent potential sites for ubiquitylation.

Using a cell based NanoBRET assay, we compared the ability of ∼100 PROTAC analogs to induce Nef ubiquitylation. Eight of the top 12 analogs identified in this screen were subsequently shown to induce Nef degradation in T cells and displayed antiretroviral activity in PBMCs. The most active analogs used thalidomide or lenalidomide as the E3 ligase targeting moiety, despite the presence of VHL ligands in 25 of the analogs used in the screen. The preference for CRBN-based degradation may result from better spatial compatibility between CRBN and Nef within the E3 ligase ternary complex. Interestingly, the NanoBRET assay also identified several analogs with z-scores significantly lower than zero, suggesting that some PROTACs may protect Nef from ubiquitylation. Ubiquitylation of both HIV-1 and SIV Nef proteins has been previously reported,^30^ raising the possibility that some PROTACs may assemble non-productive ternary complexes that interfere with endogenous ubiquitylation and turnover.

Targeted degradation of Nef may reduce HIV-1 pathogenesis and help to shrink or eliminate persistent viral reservoirs. HIV-1 gene expression, including Nef, persists in people living with HIV even in the presence of suppressive ART.^31-33^ Continued production and release of Nef and other HIV-encoded proteins from latent reservoir cells, including cells with defective proviruses,^34^ may contribute to HIV-associated co-morbidities involving the cardiovascular and central nervous systems. Furthermore, a recent study has established a significant correlation between the ability of patient-derived Nef alleles to downregulate MHC-I *in vitro* and reservoir size *in vivo*, with enhanced MHC-I downregulation activity associated with larger viral reservoirs.^20^ Our results show Nef PROTACs efficiently restore cell-surface CD4 and MHC-I expression in T cells (Figures 3 and 4), suggesting that targeted degradation is likely to reverse AP-1 and AP-2 mediated endocytosis of all cell surface receptors and restriction factors mediated by Nef. Future studies will explore the effect of Nef PROTACs on overall antiretroviral immune responses using cell-based and *in vivo* models. Furthermore, Nef PROTACs inhibit Nef-dependent enhancement of HIV-1 replication in primary cells with limited cytotoxicity (Figure 8), supporting the therapeutic potential of these Nef PROTACs.

Duette *et al*. recently reported that Nef expression is required for persistence of intact HIV-1 proviruses in effector memory T cells, a predominant reservoir cell type.^21^ Cell-surface expression of the immune check-point receptor, PD-1, is also associated with persistence of viral reservoirs *in vitro* and *in vivo*, including in the SIV/macaque model of AIDS. Keytruda (pembrolizumab) and other immune checkpoint blocking antibodies have been shown to induce latency reversal in people living with HIV^35,36^ and in SIV-infected non-human primates^37^ in the presence of suppressive ART. Importantly, Muthumani *et al*. have shown that Nef is both sufficient and necessary for HIV-induced cell-surface expression of PD-1.^38^ Taken together, these findings support a role for Nef in establishment and maintenance of the latent HIV-1 reservoir, through mechanisms involving immune escape through MHC-I downregulation, upregulation of PD-1 and possibly other immune checkpoint molecules. Targeted degradation using PROTACs is anticipated to reverse all Nef functions, potentially promoting latency reversal, restoration of adaptive immunity, suppression of viral replication and ultimately reservoir reduction.

## SIGNIFICANCE

Here we describe the synthesis and evaluation of heterobifunctional PROTAC molecules that induce the targeted degradation of the HIV-1 Nef virulence factor. Nef plays essential roles in the pathogenesis and persistence of HIV-1, and pharmacological suppression of Nef function not only inhibits viral replication but has the potential to restore the immune response to eliminate viral reservoirs. The PROTACs described here are based on previous Nef-binding compounds which display antiretroviral activity and trigger adaptive immune responses to HIV^+^ CD4 T cells *in vitro*. However, medicinal chemistry advancement of ‘occupancy based’ Nef inhibitors has been difficult because Nef lacks a singular enzymatic or biochemical activity. Instead, Nef functions by interacting with multiple host cell signaling partners through diverse structural motifs to boost viral replication and allow infected cells to hide from the immune system. The targeted degraders described here effectively block these diverse Nef functions, including downregulation of CD4 and MHC-I, which involve distinct Nef structural motifs and endolysosomal pathways. Nef PROTACs also demonstrated potent antiretroviral activity in HIV-infected primary cells. These molecules may have clinical utility in restoring immune recognition of viral reservoir cells and prevent viral rebound following traditional antiretroviral drug withdrawal. While pharmaceutical interest in PROTACs has exploded in the anticancer drug space, far fewer examples exist for infectious diseases, especially for viral accessory proteins like Nef which are generally considered undruggable. Targeted degradation of the HIV-1 Nef protein demonstrates the broader potential of antimicrobials based on PROTACs, expanding the landscape of microbial proteins for therapeutic exploitation.

## ACKNOWLEDGEMENTS

This work was supported by the National Institute of Allergy and Infectious Diseases (R01 AI152677 and R41 AI155054 to T.E.S.).

## MATERIALS AND METHODS

### PROTAC synthesis and characterization

Details of synthetic chemistry procedures and analysis of intermediates and final products are provided in the Supplemental Information.

### NanoBRET assay for PROTAC-mediated Nef ubiquitylation

Human 293T cells were cultured in DMEM supplemented with 10% fetal bovine serum. For the assay, the coding sequence of HIV-1 Nef (SF2 variant) was amplified by PCR and inserted into the plasmid pNLF-1-C (Promega) for expression of NanoLuc at the Nef C-terminus. 293T cells were plated in 6-well plates (8 x 10^5^ cells/well) and allowed to adhere for 5 h. Cells were transfected using XtremeGeneHP DNA transfection reagent (Roche) according to the manufacturer’s protocol. The pNLF-1-C-Nef plasmid (expressing Nef-nLuc) was co-transfected at a 1:5 ratio with either the HaloTag-ubiquitin plasmid (expressing Ub-Halo) or the HaloTag control (Promega). Transfected cells were incubated for 20 h at 37 °C, trypsinized, washed with PBS and resuspended in Opti-MEM reduced serum medium (Thermo Fisher/Gibco) supplemented with 4% FBS to a density of 2 x 10^5^ cells/mL. Cells were treated with either DMSO (carrier solvent; 0.1% final) or HaloTag NanoBRET 618 Ligand (Promega) and 40 µL/well were plated in a white 384-well plate (Corning #3570). PROTACs were solubilized in DMSO and added to the wells to a final concentration of 10 µM and the plates were incubated at 37 °C for 24 h. NanoBRET NanoGLO substrate (Promega) was diluted 83.3-fold in Opti-MEM without FBS and 10 µL were added to each well. The assay was read immediately on a BioTek Cytation5 plate reader using a NanoBRET fluorescence filter block with donor emission of 460 nm and acceptor emission of 618 nm. All conditions were assayed in quadruplicate, and the mean values for each well were corrected for background and expressed as 618 nm (BRET) to 460 nm (nLuc) fluorescence ratios. The ratios were normalized to the DMSO control wells and used to calculate z-scores based on the normalized ratios for all analogs tested.

### Inducible Nef expression in T cells for assessment of PROTAC activity

The CEM-T4 T cell line (HIV Reagent Program Cat. #ARP-117) was cultured in RPMI 1640 medium supplemented with 10% fetal bovine serum (FBS). Inducible expression of a Nef-eGFP fusion protein was established in CEM-T4 cells using the Lenti-XTM Tet-On System (Clontech now Takara Bio USA) which places expression of Nef-eGFP under the control of a doxycycline-inducible promoter. Following lentiviral transduction, the cells were selected with G418 (400 µg/mL) and puromycin (1 µg/mL). The selected cells were treated with doxycycline (Dox; 2 µg/mL for 48 h) and the eGFP-positive cells were sorted to obtain pure populations of Nef-eGFP cells. Cells were further sorted by CD4 staining to isolate functional Nef-eGFP which downregulates CD4 expression following Dox induction.

### Flow cytometry and immunoblotting

Inducible CEM/Nef-eGFP cells were plated at a density of 5 x 10^5^/mL in RPMI medium supplemented with 1% tetracycline-free FBS. Nef-eGFP expression was induced with doxycycline at 7.5 ng/mL (CD4 downregulation) or 75 ng/mL (MHC-I downregulation). PROTACs or DMSO vehicle as control (0.1% final) were added at the same time as doxycycline. Cells were harvested 24 h later for flow cytometry by centrifugation at 300 g for 4 min followed by 3 washes with 350 uL flow buffer (PBS supplemented with 4% FBS). Cells were resuspended in 250 uL flow buffer and stained with CD4-APC (BD Biosciences #555349) or MHC-I-PE (W6/32, Santa Cruz #sc-32235) for 45 min at room temperature in the dark. Cells were then washed 3 more times with flow buffer and resuspended in a final volume of 400 uL for flow cytometry analysis.

For immunoblotting, CEM/Nef-eGFP cells were treated with 7.5 ng/mL doxycycline for 48 h to induce Nef-eGFP expression and treated with PROTACs at the same time. Cells (2 x10^6^/sample) were collected by centrifugation at 300 g for 4 min and resuspended in cell lysis buffer (Cell Signaling #9803) supplemented with protease inhibitors (Sigma #5056489001). Cells were sonicated for 10 s and then incubated on ice for 30 minutes. Lysates were clarified by centrifugation at 15,000 g for 15 min at 4 °C, and 20 µg protein per sample was separated by SDS-PAGE and transferred to nitrocellulose membranes. Membranes were probed with HIV-1 Nef rabbit polyclonal antiserum (1:4000 dilution overnight; HIV Reagent Program) followed by an actin mouse monocolonal antibody (1:5000 dilution for 1 h; Millipore #MAB1501R). Nef and actin proteins were visualized with complementary anti-rabbit and anti-mouse secondary antibodies coupled to infrared dyes for quantitative imaging using the LI-COR platform.

### Recombinant Nef and Cereblon protein expression and purification

Expression and purification of full-length Nef proteins from multiple M-group HIV-1 variants and SIVmac239 using bacterial hosts is described in detail elsewhere.^39-41^

To create a bacterial expression vector for the human Cereblon thalidomide binding domain (TBD; amino acids 317 - 425), this coding region was PCR-amplified and subcloned via Gibson assembly into the BamHI and XhoI restriction sites of pET-28a(+)-nHis6-Smt3.^42^ The resulting coding sequence encodes for the thalidomide binding domain fused to an N-terminal Smt3 (yeast SUMO) tag with a hexa-histidine sequence at the N-terminus. Sanger DNA sequencing verified the Cereblon TBD coding sequence in the final expression plasmid.

*E. coli* strain Rosetta2(DE3)pLysS (EMD Millipore) was transformed with the pET-based Smt3-Cereblon-TBD expression plasmid. LB medium (containing 34 µg/mL chloramphenicol and 30 µg/mL kanamycin) was inoculated with a single colony and grown for 12 h at 37 °C. This starter culture was diluted 100-fold into fresh LB medium (1 L) supplemented with 20 µM ZnCl_2_, and the same antibiotics. The culture was incubated at 37 °C to an OD_600_ of 0.6 followed by cooling to 18 °C for 1h. Protein expression was induced by addition of isopropyl β-D-1-thiogalactopyranoside (IPTG) to a final concentration of 0.5 mM for 18 h. Induced cells were harvested by centrifugation, snap-frozen in liquid N_2_, and stored at −80°C.

For purification of recombinant Smt3-CereblonTBD, the cell pellet was thawed on ice and resuspended in 50 mL of Ni-IMAC (nickel-immobilized metal affinity chromatography) binding buffer [25 mM Tris-HCl, 20 mM imidazole, 500 mM NaCl, 10% (v/v) glycerol, 0.1 mM 2-mercaptoethanol (2-ME), pH 8.3]. The cell suspension was supplemented with one cOmplete, EDTA-free protease inhibitor cocktail tablet (Roche) and passed through a microfluidizer (Microfluidics) ten times at 4 °C. The lysate was clarified by centrifugation at 100,000 g for 1 h at 4 °C. The clarified lysate was loaded onto a 5 mL HisTrap HP column (Cytiva) at 2 mL/min pre-equilibrated with Ni-IMAC binding buffer. Non-specifically bound proteins were removed by a 20-column volume wash in binding buffer followed by elution with a 34-column volume linear gradient from 20 to 300 mM imidazole using Ni-IMAC elution buffer (binding buffer plus 500 mM imidazole). Fractions containing the Smt3-Cereblon-TBD were identified by SDS-PAGE, pooled, and dialyzed against 2 L of dialysis buffer (50 mM HEPES, 150 mM NaCl, 10% (v/v) glycerol, 0.1 mM 2-ME, pH 7.5) for 2 h at 4 °C, followed by a final dialysis for 12 h with fresh buffer. The His6-Smt3 tag was cleaved from the Cereblon-TBD N-terminus by addition of 150 µL of His6-Ulp1 protease (2 mg/mL) to the dialyzed protein and gentle rocking at 4 °C for 12 h. Following protease cleavage, the His6-Smt3 tag and Ulp1 protease were purified away from the Cereblon-TBD protein by application of the protein sample to a 5 mL HisTrap HP column at 2 mL/min pre-equilibrated with post-cleavage binding buffer (50 mM HEPES, 20 mM imidazole, 500 mM NaCl, 10% (v/v) glycerol, 0.1 mM 2-ME, pH 7.5). The final purification was conducted as above for the first Ni-IMAC step but used post-cleavage elution buffer (binding buffer containing 500 mM imidazole) for the elution gradient. Flow-through fractions containing the Cereblon-TBD protein were pooled, supplemented with 0.2% (w/v) n-Octyl-β-D-glucopyranoside (to prevent aggregation) and 100 µM ZnCl_2_ followed by concentration to 4.5 mL. The concentrated Cereblon-TBD protein was applied to a HiLoad 16/600 Superdex 75 size-exclusion chromatography (SEC) column (Cytiva) at 0.5 mL/min pre-equilibrated with SEC buffer (50 mM HEPES, 150 mM NaCl, 10% (v/v) glycerol, 0.1 mM TCEP, pH 7.5). SEC fractions containing pure Cereblon-TBD were pooled and concentrated to 1.1 mg/mL (93.3 µM) and frozen as aliquots at −80 °C.

### Surface plasmon resonance

SPR analysis was performed on a Reichert 4SPR instrument (Reichert Technologies) using carboxymethyl dextran hydrogel biosensor chips (Reichert #13206066). Recombinant purified Nef and CRBN proteins were covalently attached to the chip surface via standard amine coupling chemistry. PROTACs were solubilized in phosphate-buffered saline (PBS) plus 1% DMSO and injected at a flow rate of 50 µL/min for 90 s over a range of concentrations followed by a 180 s dissociation phase. The chip surface was regenerated between analogs with 5 mM NaOH at a flow rate of 50 µL/min for 30 s. Each compound was assayed in triplicate at each concentration, and the resulting sensorgrams were corrected for buffer effects and fitted with a 1:1 Langmuir binding model using TraceDrawer (Reichert). Dissociation constants were calculated from the resulting rate constants and the relationship K_D_ = k_d_/k_a_.

### PROTAC-mediated ternary complex formation in vitro

PROTAC-mediated ternary complex formation was conducted using a Superdex 75 10/300 GL analytical SEC column (Cytiva) pre-equilibrated with SEC buffer. Purified recombinant HIV-1 full-length Nef_NL4-3_ and Cereblon-TBD were thawed on ice and mixed in a 1:1 molar ratio (25 µM of each protein) in the absence or presence of a 5-fold, 10-fold, or 20-fold molar excess of Nef PROTAC. The mixtures were incubated on ice for 20 min and 100 µL was loaded onto the SEC column at a flow rate of 0.5 mL/min.

### HIV-1 replication assays

Donor peripheral blood mononuclear cells (PBMCs) were isolated from leukopaks with a PBMC isolation kit (Miltenyi Biotec). PBMCs were activated with ImmunoCult human CD3/CD28 T cell activator (STEMCELL Technologies; Cat. # 10971) and 50 U/mL IL-2 (BD-Pharmingen; Cat. # CB-40043B) for 3 days. PBMCs were infected with HIV-1_NL4-3_ for 4 d in the presence or absence of PROTACs at a final concentration of 1.0 µM with DMSO as carrier solvent (0.1% final concentration). Parallel cultures were infected with Nef-defective HIV-1 to assess the extent of Nef-mediated enhancement of replication. Cells were lysed with 0.1% Triton X-100 and viral titers measured by AlphaLISA assay (PerkinElmer) for HIV-1 p24. Cell viability was assessed in uninfected PBMC incubated with each PROTAC under the same conditions (1.0 µM, 4 d) using the CellTiter-Blue assay (Promega).

