## Supplemental Information for "PROTAC-mediated Degradation of HIV-1 Nef Efficiently Restores Cell-surface CD4 and MHC-I Expression and Blocks HIV-1 Replication"

**Figure S1**. NanoBRET ubiquitylation assay data for active Nef PROTACs.

**Figure S2.** Chemical structures of analogs with little to no activity in the receptor downregulation and Nef degradation assays.

**Figure S3.** Nef PROTACs stabilize ternary complexes of recombinant purified Nef with the thalidomide-binding domain of CRBN *in vitro*.

**Figure S4**. Comparison of HIV-1 Nef (B-clade variant NL4-3) and SIV Nef (mac239) crystal structures and primary sequences.

**Table S1.** Surface Plasmon Resonance (SPR) analysis of thalidomide analog binding to Nef proteins and to the Cereblon thalidomide-binding domain (TBD).

**Synthetic chemistry:** Detailed synthetic methods and analytical chemistry of Nef PROTAC analogs.

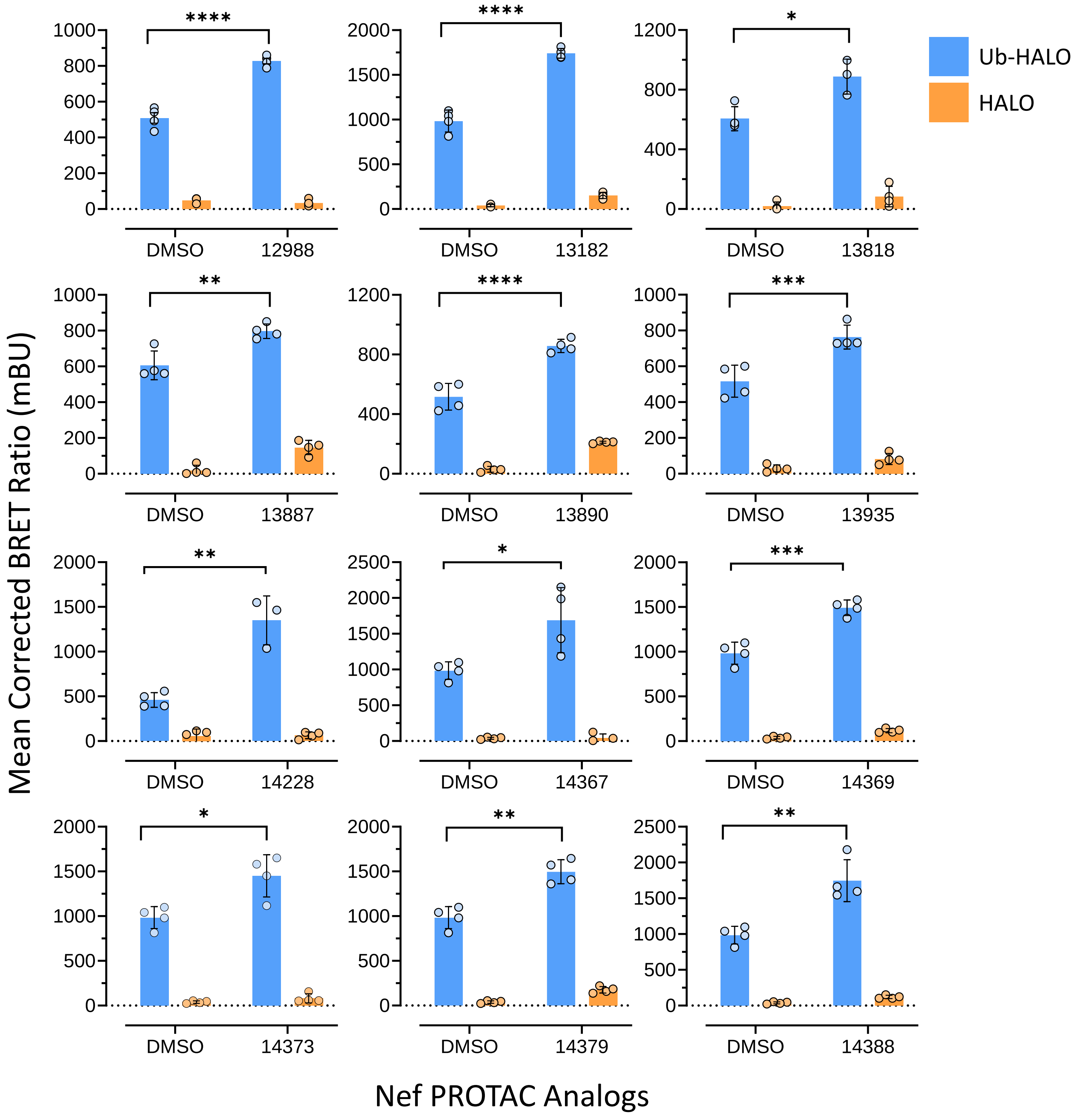


**Figure S1. NanoBRET ubiquitylation assay data for active Nef PROTACs.** In this assay, Nef is fused to nano-Luciferase (Nef-nLuc) and co-expressed with a ubiquitin-Halo tag fusion protein (Ub-Halo) in 293T cells. PROTACs promote ligation of Ub-Halo to Nef-nLuc, which is detected by bioluminescence resonance energy transfer (BRET) from nLuc to the Halo Tag (see main Figure 2). Each compound was assayed in quadruplicate (individual points shown) and the average 618 nm to 460 nm fluorescence ratios (BRET signal for Ub incorporation normalized to Nef-nLuc levels) are represented as the blue bar heights ± SD. Basal ubiquitylation of Nef was determined in the absence of each PROTAC (DMSO control). Quadruplicate control reactions were run in parallel with the Halo tag alone (orange bars) and the resulting Halo-only BRET ratios were significantly lower than the values obtained with Ub-Halo in each case (ranging from 4% to 25% of the specific signals). All twelve active PROTACs significantly increased the Ub-dependent BRET ratio compared to the DMSO controls (Student's t test; *, $p < 0.05$; **, $p < 0.01$; ***, $p < 0.001$; ****, $p < 0.0001$).

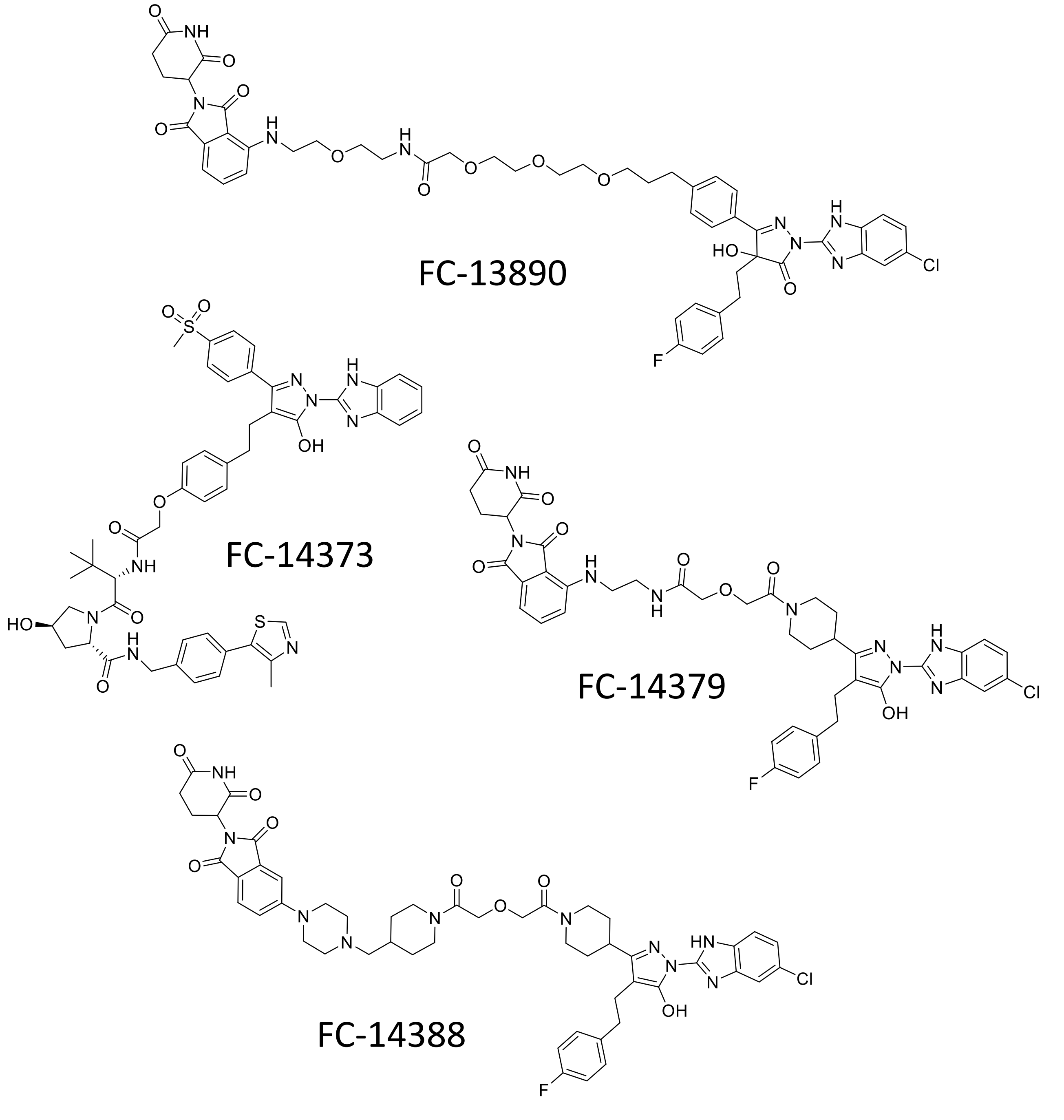


**Figure S2. Chemical structures of analogs with little to no activity in the receptor downregulation and Nef degradation assays.** Data with these analogs are referenced in Figures 3 and 4 in the main text.

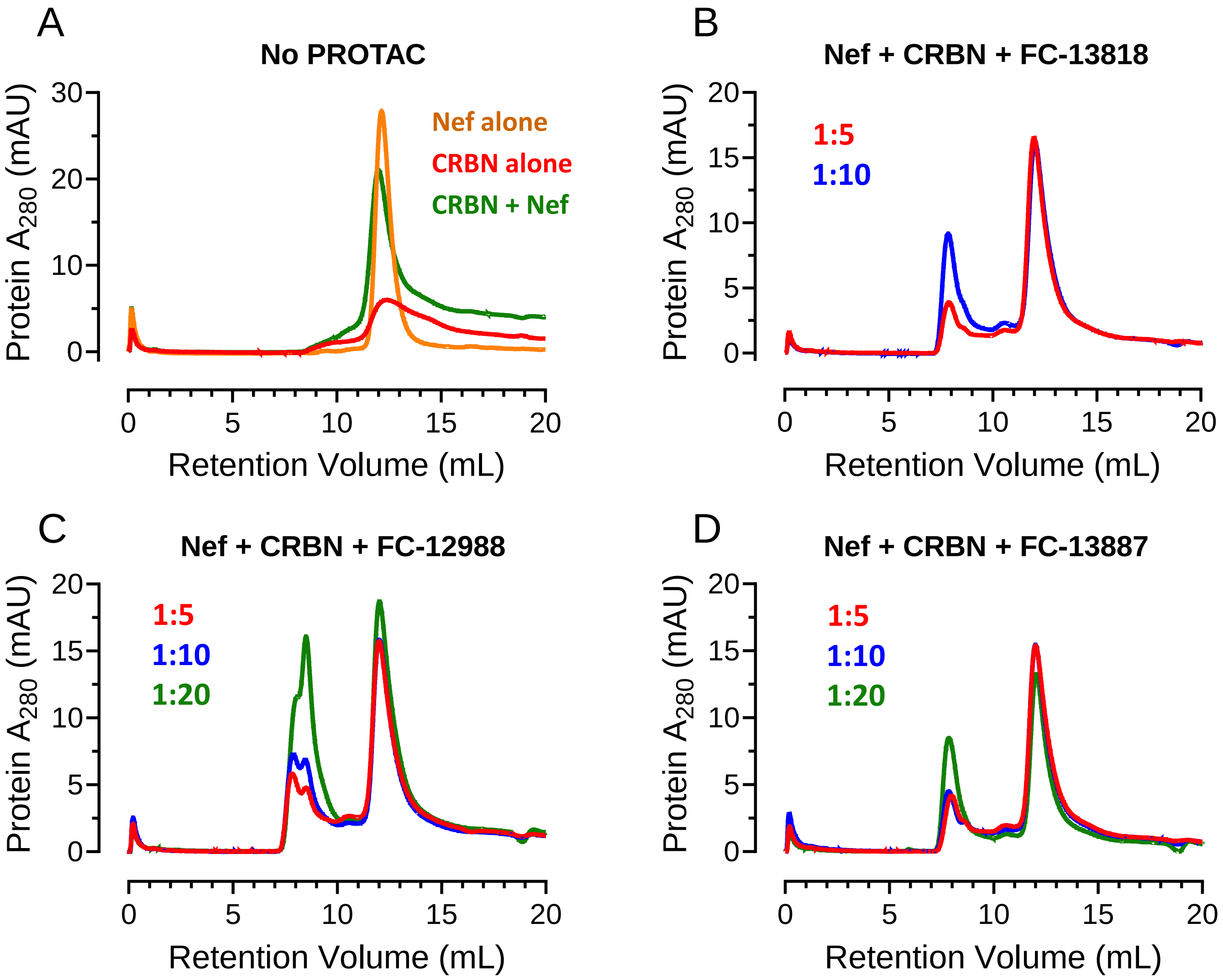


**Figure S3. Nef PROTACs stabilize ternary complexes of recombinant purified Nef with the thalidomide-binding domain of CRBN *in vitro*.** A) Recombinant purified HIV-1 Nef (NL4-3 variant) and the CRBN ligand-binding domain were analyzed by size-exclusion chromatography (Superdex 75 column) either individually or after mixing in equimolar proportions. Note that the mixture elutes as a single peak as the individual proteins have very similar retention volumes. B-C) The Nef-CRBN protein mixture from part A was incubated with Nef PROTACs **FC-13818**, **12988**, and **13887** in the molar ratios shown for 20 min at 4 °C prior to SEC. The new peaks with smaller retention volumes (higher molecular weights) correspond to ternary complexes of Nef, CRBN, and the PROTAC.

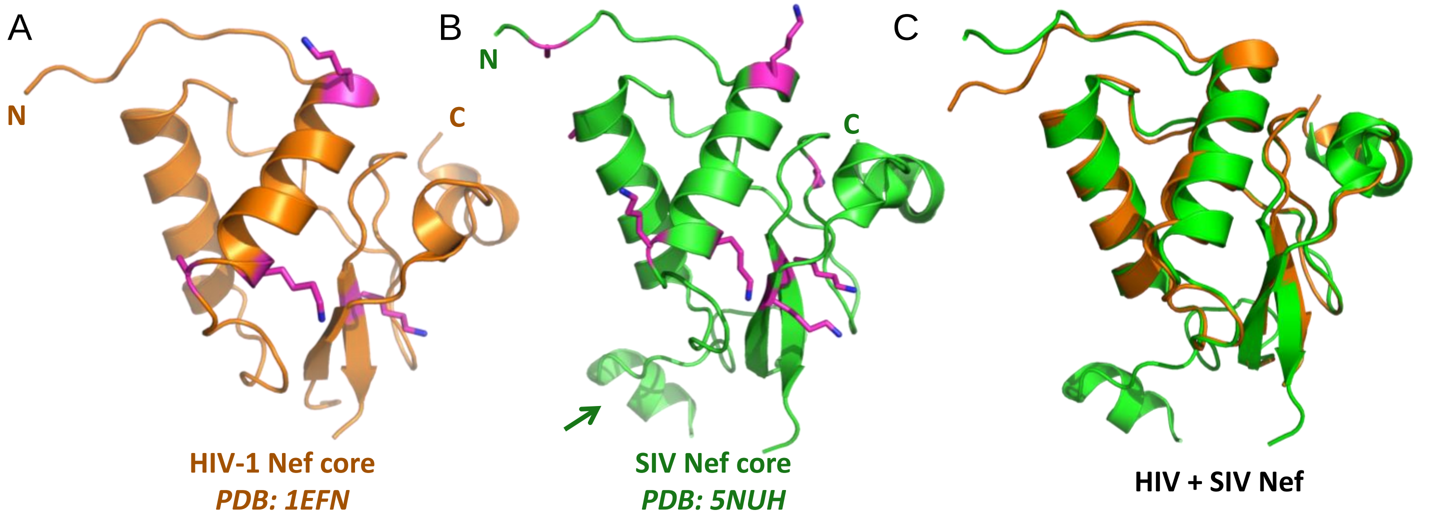


D

```
HIV: MGGKWSKRSM GGWSAIRERM RRAEPRAEP- ----AADG-- --VGAVSRDL EKHGAITSSN TAATNADCAW LEAQEEEE-- ----------
SIV: MGGAISMRRS RPSGDLRQRL LRARGETYGR LLGEVEDGYS QSPGGLDKGL SSLSCEGQKY NQGQYMNTPW RNPAEEREKL AYRKQNMDDI

HIV: -------VGF PVRPQVPLRP MTYKAALDIS HFLKEKGGLE GLIWSQRRQE ILDLWIYHTQ GYFPDWQNYT PGPGIRYPLT FGWCFKLVPV
SIV: DEEDDDLVGV SVRPKVPLRT MSYKLAIDMS HFIKEKGGLE GIYYSARRHR ILDIYLEKEE GIIPDWQDYT SGPGIRYPKT FGWLWKLVPV

HIV: EPEKVEEANE GENNSLLHPM SLHGMEDAEK EVLVWRFDSK LAFHHMARELH PEYFKDC-- ---------- ---------- -----
SIV: NVS--DEAQE DEEHYLMHPA QTSQWDDPWG EVLAWKFDPT LAYTYEAYVRY PEEFGSKSG LSEEEVRRRL TARGLLNMAD KKETR
```

**Figure S4. Comparison of HIV-1 Nef$_{NL4\text{-}3}$ and SIV$_{mac239}$ Nef structures and primary sequences.** Models were produced using PyMol and the crystal coordinates indicated. A) HIV-1 Nef and B) SIV Nef core regions are rendered in orange and green respectively, with lysine residues (potential Ub sites) highlighted in magenta. The N- and C-terminal ends of each protein are also indicated. Unstructured regions including the N-terminal anchor domain and an internal loop were not present in the electron density. C) Structural alignment of SIV and HIV Nef core domains shows a very similar overall core fold. D) Amino acid sequence alignment of the complete SIV and HIV Nef proteins used for crystallography. Identical and homologous residues are highlighted in red. The core regions present in the structural models are underlined. Note that a portion of the SIV Nef internal loop forms a short α-helix in SIV Nef (*arrow* in panel B); this region is unstructured in HIV-1 Nef in the absence of a binding partner.

**Table S1. Surface Plasmon Resonance (SPR) analysis of thalidomide analog binding to Nef proteins and to the Cereblon thalidomide binding domain (TBD).** The recombinant purified Nef and CRBN-TBD proteins shown were covalently attached to carboxymethyl dextran chips via standard amine coupling chemistry. Compounds were solubilized in phosphate-buffered saline plus 1% DMSO and injected at a flow rate of 50 µL/min for 90 s over a range of concentrations followed by a 180 s dissociation phase. The chip surface was regenerated between analogs with 5 mM NaOH at a flow rate of 50 µL/min for 30 s. Each compound was assayed in triplicate at each concentration, and the resulting sensorgrams were corrected for buffer effects and fitted with a 1:1 Langmuir binding model using TraceDrawer (Reichert). Dissociation constants were calculated from the resulting rate constants and the relationship $K_D = k_d/k_a$. Structures of the compounds tested are shown below the table.

| | **$K_D$ values, M** | | | |
|---|---|---|---|---|
| **Compound** | **HIV-1 Nef*SF2*** | **HIV-1 Nef*NL4-3*** | **SIV Nef*mac239*** | **CRBN** |
| **FC-13454** | $2.20 \times 10^{-5}$ | $1.24 \times 10^{-5}$ | $2.18 \times 10^{-5}$ | $1.85 \times 10^{-6}$ |
| **FC-13455** | $6.34 \times 10^{-5}$ | $2.48 \times 10^{-5}$ | $6.55 \times 10^{-5}$ | $4.27 \times 10^{-6}$ |
| **FC-13457** | $4.41 \times 10^{-5}$ | $2.93 \times 10^{-5}$ | $2.28 \times 10^{-5}$ | $1.99 \times 10^{-6}$ |
| **FC-13458** | $3.23 \times 10^{-5}$ | $2.97 \times 10^{-5}$ | $3.80 \times 10^{-5}$ | $3.64 \times 10^{-6}$ |
| **FC-13459** | $1.69 \times 10^{-5}$ | $1.35 \times 10^{-5}$ | $1.38 \times 10^{-5}$ | $7.87 \times 10^{-6}$ |
| **FC-13460** | $2.74 \times 10^{-5}$ | $1.86 \times 10^{-5}$ | $3.35 \times 10^{-5}$ | $1.32 \times 10^{-6}$ |

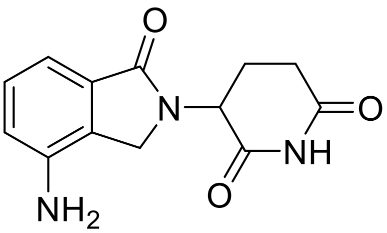

**FC-13454**
(lenalidomide)

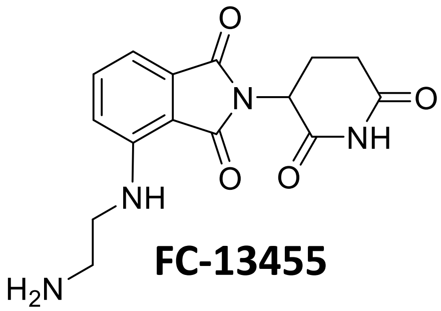

**FC-13455**

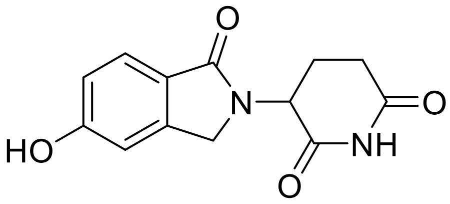

**FC-13457**

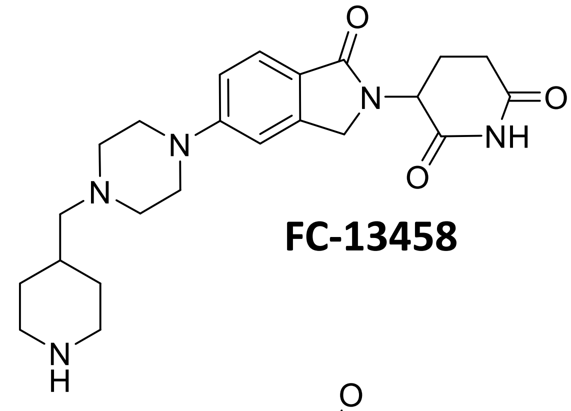

**FC-13458**

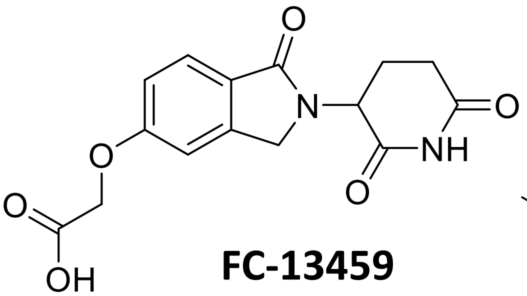

**FC-13459**

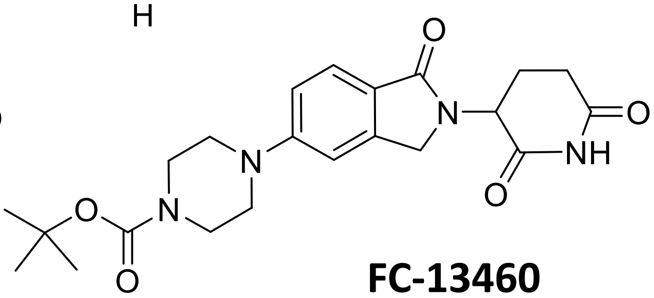

**FC-13460**

**Synthetic chemistry:**

**General experimental methods**

Starting reagents were purchased from commercial suppliers and were used without further purification unless otherwise specified. Normal phase column chromatography was carried out in the indicated solvent system using pre-packed silica gel cartridges on the Isco CombiFlash Companion® or Isco CombiFlash Rf® system. Preparative reverse phase HPLC ("Prep HPLC") was performed on a Phenomenex LUNA 5 µm C18(2) 100 Å 150 x 21.2 mm column or a Waters SunFire® Prep C18 OBD™ 10 µm 150 x 30 mm column with a 12 min mobile phase gradient of 10% acetonitrile/water to 90% acetonitrile/water with 0.1% TFA as buffer using 214 and 254 nm as detection wavelengths. Injection and fraction collection were performed with a Gilson 215 liquid handling apparatus using Unipoint software.

LC-MS data were determined with a Waters Alliance 2695 HPLC/MS using a Phenomenex Luna 3 µm C18(2) 100 Å, 75 x 4.6 mm column with a 2996 diode array detector operating from 210–400 nm; the solvent system was 5–95% acetonitrile in water (with 0.1% TFA) over nine minutes using a linear gradient, and retention times ($t_R$) are in minutes. Mass spectrometry was performed on a Waters ZQ using electrospray in positive ion mode. High resolution mass spectra (HRMS) were obtained on a Bruker Daltonics microTOF II instrument using electrospray ionization in positive mode.

Nuclear Magnetic Resonance spectra were recorded on a Varian Mercury 300 spectrometer operating at 299.985 MHz for $^1$H NMR, at 282.243 MHz for $^{19}$F NMR and at 75.439 MHz for $^{13}$C NMR. Spectra were taken in the indicated solvent at ambient temperature, and the chemical shifts are reported in parts per million (ppm, δ) relative to the lock of the solvent used. Resonance patterns are recorded with the following notations: br (broad), s (singlet), d (doublet), t (triplet), q (quartet), and m (multiplet).

**Synthesis of CRBN binding intermediates**

**2-{[2-(2,6-dioxopiperidin-3-yl)-1-oxo-2,3-dihydro-1H-isoindol-4-yl]oxy}acetic acid**

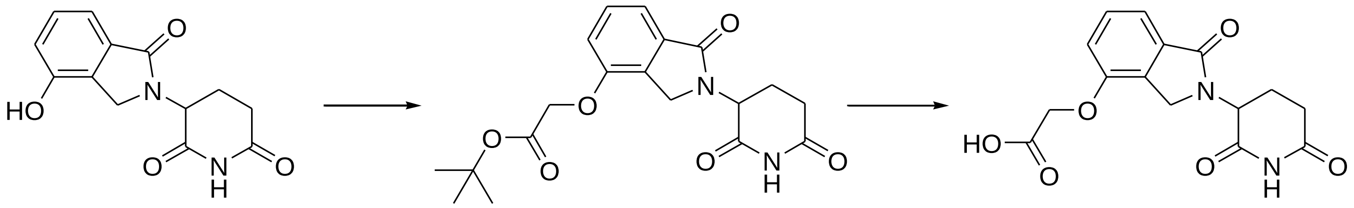

*tert-butyl 2-{[2-(2,6-dioxopiperidin-3-yl)-1-oxo-2,3-dihydro-1H-isoindol-4-yl]oxy}acetate*. To a stirred solution of 3-(4-hydroxy-1-oxo-2,3-dihydro-1H-isoindol-2-yl)piperidine-2,6-dione (926 mg, 3.6 mmol) and i-$Pr_2NEt$ (0.64 mL, 3.6 mmol) in dry DMF (10 mL) was added t-butyl bromoacetate (0.5 mL, 3.4 mmol). The mixture was stirred at 40 °C for 2 d and purified directly by prep HPLC to give the title compound (380 mg, 28%) as an oil.

$^1$H NMR (300MHz, DMSO-$d_6$) δ = 7.52 - 7.39 (m, 1H), 7.32 (d, *J*=7.0 Hz, 1H), 7.21 - 7.08 (m, 1H), 5.10 (dd, *J*=5.1, 13.0 Hz, 1H), 4.82 (s, 2H), 4.48 - 4.13 (m, 2H), 3.03 - 2.75 (m, 1H), 2.63-2.33 (m, 2H), 2.14 - 1.88 (m, 1H), 1.40 (s, 9H).

LC-MS $t_R$ = 3.90 min, m/z 375, 319.

*2-{[2-(2,6-dioxopiperidin-3-yl)-1-oxo-2,3-dihydro-1H-isoindol-4-yl]oxy}acetic acid.* A solution of tert-butyl 2-{[2-(2,6-dioxopiperidin-3-yl)-1-oxo-2,3-dihydro-1H-isoindol-4-yl]oxy}acetate (380 mg, 1.0 mmol) in 2:1 $CH_2Cl_2$/TFA (9 mL) was stirred at rt for 1 d and concentrated. The residue was lyophilized from aq MeCN to give the title compound (289 mg, 90%) as an off-white solid.

$^1$H NMR (300MHz, DMSO-d6) δ = 7.52 - 7.40 (m, 1H), 7.32 (d, *J*=7.5 Hz, 1H), 7.14 (d, *J*=7.9 Hz, 1H), 5.10 (d, *J*=8.3 Hz, 1H), 4.82 (s, 2H), 4.45 - 4.15 (m, 2H), 3.02 - 2.78 (m, 1H), 2.64-2.20 (m, 2H), 2.10 - 1.86 (m, 1H).

**4-[(2-Aminoethyl)amino]-2-(2,6-dioxopiperidin-3-yl)-2,3-dihydro-1H-isoindole-1,3-dione**

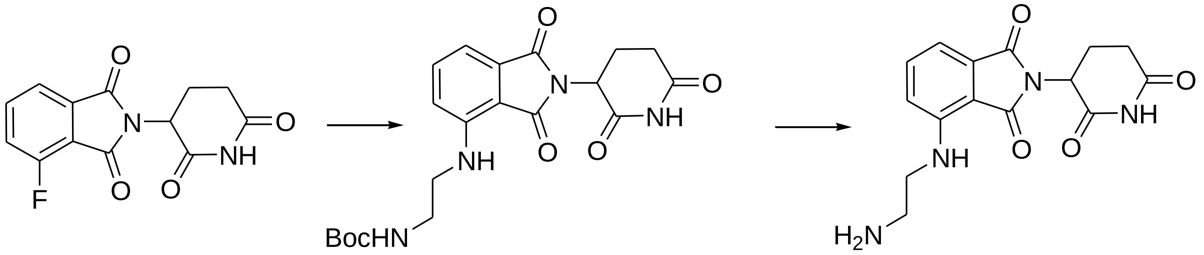


*tert-Butyl N-(2-{[2-(2,6-dioxopiperidin-3-yl)-1,3-dioxo-2,3-dihydro-1H-isoindol-4-yl]amino}ethyl)carbamate*. A mixture of 2-(2,6-dioxopiperidin-3-yl)-4-fluoro-2,3-dihydro-1H-isoindole-1,3-dione (1.75 g, 6.3 mmol), tert-butyl N-(2-aminoethyl)carbamate (1.06 g, 6.6 mmol), i-$Pr_2NEt$ (2.3 mL, 12.7 mmol) and dry DMF (30 mL) was stirred in a 90°C oil bath for 2 d. The mixture was diluted with EtOAc (150 mL) and washed with water (2 x 50 mL) and brine (50 mL). The combined aqueous washes were back extracted with EtOAc (50 mL). The combined EtOAc layer was dried over $Na_2SO_4$ and concentrated under reduced pressure to leave a black oil (4.11 g). Prep HPLC gave the compound (850 mg, 32%) as a yellow solid.

$^1$H NMR (300MHz, DMSO-$d_6$) Shift = 7.64 - 7.42 (m, 1H), 7.10 (d, J=8.8 Hz, 1H), 7.04 - 6.93 (m, 2H), 6.68 (br s, 1H), 5.01 (dd, J=5.5, 12.5 Hz, 1H), 3.38-3.21 (m, 2H), 3.14-3.01 (m, 2H), 2.92 - 2.68 (m, 1H), 2.59-2.32 (m, 2H), 2.11 - 1.80 (m, 1H), 1.32 (s, 9H).

LC-MS $t_R$ 3.98 min, m/z 439, 417, 317.

*4-[(2-Aminoethyl)amino]-2-(2,6-dioxopiperidin-3-yl)-2,3-dihydro-1H-isoindole-1,3-dione*. A solution of tert-butyl N-(2-{[2-(2,6-dioxopiperidin-3-yl)-1,3-dioxo-2,3-dihydro-1H-isoindol-4-yl]amino}ethyl)carbamate (850 mg, 2.0 mmol) in 2:1 $CH_2Cl_2$/TFA (30 mL) was stirred at rt for 0.5 h. The mixture was concentrated, and the residue was lyophilized from MeCN/5% aq HCl to give the title compound (745 mg, quant) as its HCl salt.

$^1$H NMR (300MHz, $CD_3OD$) Shift = 7.62 (t, J=7.7 Hz, 1H), 7.15 (d, J=8.3 Hz, 2H), 5.16 - 5.01 (m, 1H), 3.68 (s, 2H), 3.26 - 3.10 (m, 2H), 2.95 - 2.59 (m, 3H), 2.23 - 2.00 (m, 1H).

LC-MS $t_R$ 2.25 min, m/z 317.

**4-[(2-aminoethyl)(methyl)amino]-2-(2,6-dioxopiperidin-3-yl)-2,3-dihydro-1H-isoindole-1,3-dione**

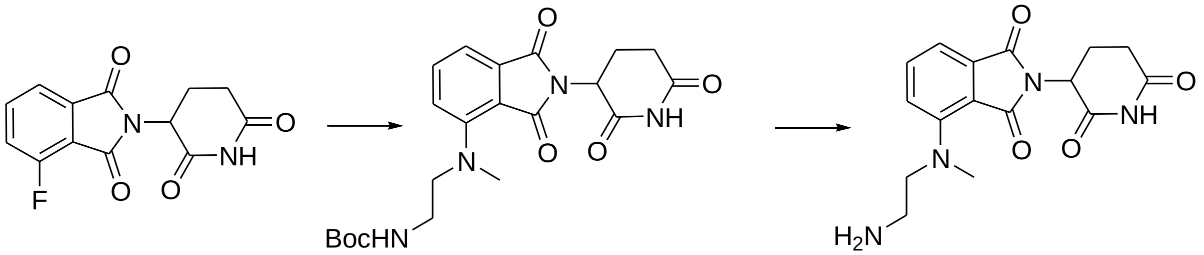


*tert-butyl N-(2-{[2-(2,6-dioxopiperidin-3-yl)-1,3-dioxo-2,3-dihydro-1H-isoindol-4-yl](methyl)amino}ethyl)carbamate*. A mixture of 2-(2,6-dioxopiperidin-3-yl)-4-fluoro-2,3-dihydro-1H-isoindole-1,3-dione (750 mg, 2.7 mmol), tert-butyl N-[2-(methylamino)ethyl]carbamate (500 mg, 2.9 mmol), i-$Pr_2NEt$ (1 mL, 5.5 mmol) and DMF (10 mL) was stirred at 70 °C for 20 h. The mixture was cooled to rt, diluted with EtOAc (90 mL), washed with water (2 x 20 mL) and brine (20 mL), and dried over $Na_2SO_4$. Removal of the solvent left a yellow oil (1.58 g). Prep HPLC gave the title compound (480 mg, 41%) as a yellow solid.

$^1$H NMR (300MHz, DMSO-$d_6$) δ = 7.82 - 7.67 (m, 1H), 7.46 - 7.30 (m, 2H), 6.94 - 6.79 (m, 1H), 5.32 - 5.17 (m, 1H), 3.81 - 3.57 (m, 2H), 3.35 - 3.22 (m, 2H), 3.12 - 2.93 (m, 1H), 2.81 - 2.60 (m, 2H), 2.24 - 2.02 (m, 1H), 1.42 (s, 9H).

*4-[(2-aminoethyl)(methyl)amino]-2-(2,6-dioxopiperidin-3-yl)-2,3-dihydro-1H-isoindole-1,3-dione.* A solution of tert-butyl N-(2-{[2-(2,6-dioxopiperidin-3-yl)-1,3-dioxo-2,3-dihydro-1H-isoindol-4-yl](methyl)amino}ethyl)carbamate (480 mg, 1.1 mmol) in 2:1 $CH_2Cl_2$/TFA (30 mL) was stirred at rt for 0.5 h and concentrated. The residue was lyophilized from a mixture of MeCN and 5% aq HCl to give the HCl salt of the title compound (419 mg, quant) as a yellow solid.

$^1$H NMR (300MHz, $CD_3OD$) δ = 7.80 - 7.66 (m, 1H), 7.53 - 7.35 (m, 2H), 5.20-5.10 (m, 1H), 3.65-3.57 (m, 2H), 3.41 - 3.30 (m, 2H), 2.98 (s, 3H), 2.94 - 2.58 (m, 3H), 2.21 - 2.06 (m, 1H).

**4-({2-[2-(2-aminoethoxy)ethoxy]ethyl}amino)-2-(2,6-dioxopiperidin-3-yl)-2,3-dihydro-1H-isoindole-1,3-dione**

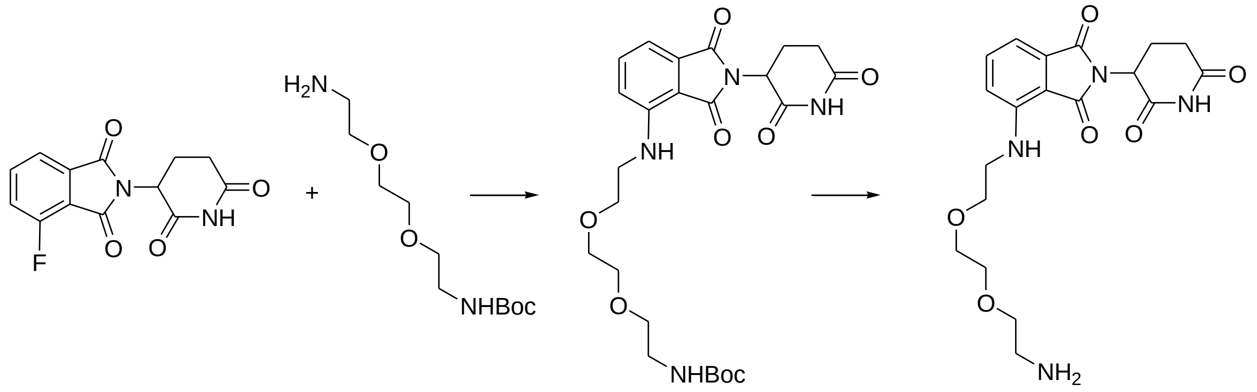

*tert-butyl N-{2-[2-(2-{[2-(2,6-dioxopiperidin-3-yl)-1,3-dioxo-2,3-dihydro-1H-isoindol-4-yl]amino}ethoxy)ethoxy]ethyl}carbamate*. A solution of 2-(2,6-dioxopiperidin-3-yl)-4-fluoro-2,3-dihydro-1H-isoindole-1,3-dione (420 mg, 1.5 mmol), tert-butyl N-{2-[2-(2-aminoethoxy)ethoxy]ethyl}carbamate (396 mg, 1.6 mmol) and i-$Pr_2NEt$ (0.55 mL, 3.0 mmol) in DMF (10 mL) was stirred at 70 °C for 4 h. The mixture was diluted with EtOAc (90 mL), washed with 5% aq HCl (2 x 15 mL) and brine (15 mL), and dried over $Na_2SO_4$. Removal of the solvent left a green oil (910 mg) which was purified by prep HPLC to give the title compound (230 mg, 30%) as a yellow solid.

$^1H$ NMR (300MHz, DMSO-$d_6$) δ = 7.62 - 7.49 (m, 1H), 7.13 (d, *J*=8.8 Hz, 1H), 7.02 (d, *J*=7.0 Hz, 1H), 6.81 - 6.67 (m, 1H), 6.65 - 6.55 (m, 1H), 5.10 - 4.98 (m, 1H), 3.67 - 3.41 (m, 10H), 3.39-3.32 (m, 2H), 3.11 - 2.97 (m, 2H), 2.95 - 2.74 (m, 1H), 2.65 - 2.36 (m, 2H), 2.08 - 1.92 (m, 1H), 1.34 (s, 9H). LC-MS $t_R$ = 4.13 min, m/z 505, 405.

*4-({2-[2-(2-aminoethoxy)ethoxy]ethyl}amino)-2-(2,6-dioxopiperidin-3-yl)-2,3-dihydro-1H-isoindole-1,3-dione.* A solution of tert-butyl N-{2-[2-(2-{[2-(2,6-dioxopiperidin-3-yl)-1,3-dioxo-2,3-dihydro-1H-isoindol-4-yl]amino}ethoxy)ethoxy]ethyl}carbamate (230 mg, 0.46 mmol) in 3:2 $CH_2Cl_2$/TFA (5 mL) was stirred at rt for 0.5 h and concentrated. The residue was lyophilized from a mixture of MeCN and 5% aq HCl to give the HCl salt of the title compound (114 mg, 57%) as a greenish yellow solid.

$^1H$ NMR (300MHz, DMSO-$d_6$) δ = 8.06 (br s, 2H), 7.65 - 7.50 (m, 1H), 7.14 (d, *J*=8.8 Hz, 1H), 7.03 (d, *J*=7.0 Hz, 1H), 6.00 - 5.21 (m, 2H), 5.12-5.00 (m, 1H), 3.70 - 3.52 (m, 8H), 3.50-3.40 (m, 2H), 2.99-2.80 (m, 2H), 2.66 - 2.37 (m, 1H), 2.10 - 1.90 (m, 1H)

**2-(2,6-dioxopiperidin-3-yl)-5-{4-[(piperidin-4-yl)methyl]piperazin-1-yl}-2,3-dihydro-1H-isoindole-1,3-dione**

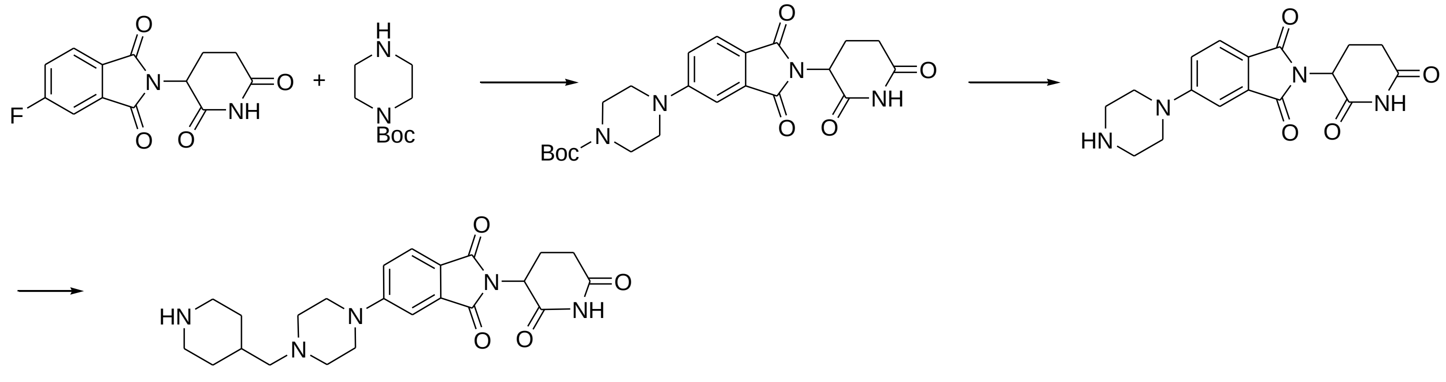

*tert-butyl 4-[2-(2,6-dioxopiperidin-3-yl)-1,3-dioxo-2,3-dihydro-1H-isoindol-5-yl]piperazine-1-carboxylate*. To a stirred solution of 2-(2,6-dioxopiperidin-3-yl)-5-fluoro-2,3-dihydro-1H-isoindole-1,3-dione (800 mg, 2.9 mmol) and tert-butyl piperazine-1-carboxylate (560 mg, 3.5 mmol) in DMSO (10 mL) was added i-$Pr_2NEt$ (1.1 mL, 6.1 mmol). The mixture was stirred at 90 °C for 3 h, cooled to rt and purified by prep HPLC to give the title compound (410 mg, 32%) as a yellow solid.

$^1H$ NMR (300MHz, DMSO-$d_6$) Shift = 7.74 - 7.64 (m, 1H), 7.33 (d, J=2.2 Hz, 1H), 7.23 (d, J=8.8 Hz, 1H), 5.06 (dd, J=5.5, 12.5 Hz, 1H), 3.45 (s, 8H), 2.99 - 2.75 (m, 1H), 2.63 - 2.51 (m, 2H), 2.08 - 1.89 (m, 1H), 1.41 (s, 9H). LC-MS $t_R$ = 4.28 min, m/z 386, 342.

*2-(2,6-dioxopiperidin-3-yl)-5-(piperazin-1-yl)-2,3-dihydro-1H-isoindole-1,3-dione*. To a stirred solution of tert-butyl 4-[2-(2,6-dioxopiperidin-3-yl)-1,3-dioxo-2,3-dihydro-1H-isoindol-5-yl]piperazine-1-carboxylate (410 mg, 0.93 mmol) in $CH_2Cl_2$ (6 mL) was added TFA (2 mL). The mixture was stirred at rt for 1 h and concentrated. The residue was lyophilized from MeCN/5% aq HCl to give the bis HCl salt of the title compound (450 mg, quant) as a yellow solid.

LC-MS $t_R$ = 2.23 min, m/z 342.

*tert-butyl 4-({4-[2-(2,6-dioxopiperidin-3-yl)-1,3-dioxo-2,3-dihydro-1H-isoindol-5-yl]piperazin-1-yl}methyl)piperidine-1-carboxylate*. To a stirred mixture of the bis HCl salt of 2-(2,6-dioxopiperidin-3-yl)-5-(piperazin-1-yl)-2,3-dihydro-1H-isoindole-1,3-dione (450 mg, 1.1 mmol), tert-butyl 4-formylpiperidine-1-carboxylate (460 mg, 2.1 mmol), NaOAc (270 mg, 3.3 mmol) and dry DCE (10 mL) was added $MgSO_4$ (150 mg). The mixture was stirred under $N_2$ for 0.5 h and $NaBH(OAc)_3$ (689 mg, 3.3 mmol) was added. The mixture was stirred overnight at rt, diluted with water (10 mL) and concentrated under reduced pressure to remove DCE. The residue was diluted with saturated aqueous $NaHCO_3$ (20 mL) and extracted with EtOAc (2 x 80 mL). The combined EtOAc layer was concentrated to leave a yellow solid (680 mg) which was purified by prep HPLC to give the TFA salt of the title compound (320 mg, 45%).

$^1$H NMR (300MHz, DMSO-$d_6$) δ = 7.81 - 7.71 (m, 1H), 7.54 - 7.45 (m, 1H), 7.41 - 7.24 (m, 1H), 5.17 - 4.98 (m, 1H), 4.31 - 4.09 (m, 2H), 4.03 - 3.80 (m, 2H), 3.65-3.05 (m, 10H), 2.95-2.40 (m, 4H), 2.10 - 1.89 (m, 2H), 1.82 - 1.62 (m, 2H), 1.38 (s, 9H), 1.17 - 0.93 (m, 1H).

LC-MS $t_R$ = 3.25 min, m/z 540, 484, 440.

*2-(2,6-dioxopiperidin-3-yl)-5-{4-[(piperidin-4-yl)methyl]piperazin-1-yl}-2,3-dihydro-1H-isoindole-1,3-dione.* A solution of the TFA salt of tert-butyl 4-({4-[2-(2,6-dioxopiperidin-3-yl)-1,3-dioxo-2,3-dihydro-1H-isoindol-5-yl]piperazin-1-yl}methyl)piperidine-1-carboxylate (320 mg, 0.49 mmol) in 3:1 $CH_2Cl_2$/TFA (6 mL) was stirred at rt for 1 h and concentrated. The residue was lyophilized from MeCN/5% aq HCl to give the bis HCl salt of the title compound (306 mg, quant) as a solid.

LC-MS $t_R$ = 2.16 min, m/z 440.

**Synthesis of Nef binding intermediates**

**2-(4-{2-[1-(5-chloro-1H-1,3-benzodiazol-2-yl)-5-hydroxy-3-[4-(trifluoromethyl)phenyl]-1H-pyrazol-4-yl]ethyl}phenoxy)acetic acid**

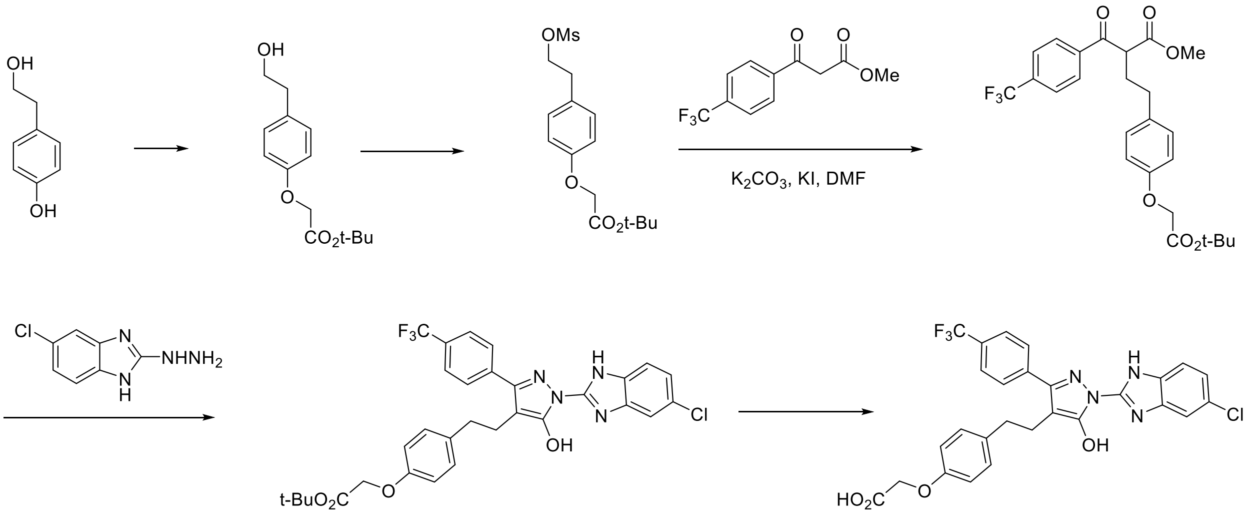


*tert-Butyl 2-[4-(2-hydroxyethyl)phenoxy]acetate*. To a stirred mixture 4-(2-hydroxyethyl)phenol (4.97 g, 36.0 mmol), $K_2CO_3$ (5.47 g, 39.6 mmol) and DMF (50 mL) was added t-butyl bromoacetate (5.3 mL, 36.0 mmol). The mixture was stirred at rt for 1 d, diluted with water (150 mL) and extracted with EtOAc (4 x 50 mL). The combined EtOAc layer was washed with water (25 mL) and brine (25 mL), and dried over $Na_2SO_4$. Removal of the solvent left a viscous oil (13.20 g) which was purified by chromatography on an 80 g silica cartridge, eluted with a 0-70% EtOAc in hexanes gradient to provide the title compound (7.28 g, 80%) as a colorless oil.

$^1$H NMR (300MHz, $CDCl_3$) δ = 7.13 (d, *J*=8.8 Hz, 2H), 6.84 (d, *J*=8.4 Hz, 2H), 4.49 (s, 2H), 3.89 - 3.69 (m, 2H), 2.87 - 2.65 (m, 2H), 1.45 (s, 9H).

*tert-butyl 2-{4-[2-(methanesulfonyloxy)ethyl]phenoxy}acetate*. To a stirred, ice-cold solution of tert-butyl 2-[4-(2-hydroxyethyl)phenoxy]acetate (7.28 g, 28.9 mmol) and i-$Pr_2NEt$ (15.5 mL, 86.6 mmol) in $CH_2Cl_2$ (150 mL) was added methanesulfonyl chloride (5.6 mL, 72.2 mmol). The ice bath was allowed to melt, and the mixture was stirred overnight at rt. The mixture was concentrated, and the residue was taken up in EtOAc (150 mL), washed with 5% aq HCl (2 x 50 mL) and brine (50 mL). The EtOAc layer was filtered through Celite. The filtrate was dried over $Na_2SO_4$ and concentrated to leave the title compound (9.72 g, quant.) as a dark red oil.

$^1$H NMR (300MHz, $CDCl_3$) δ = 7.11 (m, 2H), 6.88 - 6.76 (m, 2H), 4.47 (s, 2H), 4.40 - 4.28 (m, 2H), 3.02 - 2.91 (m, 2H), 2.81 (s, 3H), 1.46 (s, 9H).

*Methyl 4-{4-[2-(tert-butoxy)-2-oxoethoxy]phenyl}-2-[4-(trifluoromethyl)benzoyl]butanoate*. A mixture of methyl 3-oxo-3-[4-(trifluoromethyl)phenyl]propanoate (5.00 g, 20.3 mmol), tert-butyl 2-{4-[2-(methanesulfonyloxy)ethyl]phenoxy}acetate (6.84 g, 20.7 mmol), $K_2CO_3$ (2.81 g, 20.3 mmol), KI (3.37 g, 20.3 mmol) and DMF (50 mL) was stirred at 70 °C for 6 h. The mixture was cooled, diluted with EtOAc (350 mL), washed with 5% aq HCl (50 mL), water (50 mL) and brine (50 mL), and dried over $Na_2SO_4$. Removal of the solvent left a dark brown oil (12.06 g) which was chromatographed on a 120 g silica cartridge, eluted with a 0-25% EtOAc in hexanes gradient, to provide a ~3:1 mixture of desired title compound and O-alkylation product methyl (2)-3-(2-{4-[2-(tert-butoxy)-2-oxoethoxy]phenyl}ethoxy)-3-[4-(trifluoromethyl)phenyl]prop-2-enoate (4.89 g, 50%) as a pale yellow oil.

$^1$H NMR (300MHz, $CDCl_3$) δ = 7.99 - 7.88 (m, 2H), 7.78 - 7.65 (m, 2H), 7.11 - 7.02 (m, 2H), 6.88 - 6.74 (m, 2H), 4.54 - 4.44 (s, 2H), 3.64 (s, 3H), 3.12 - 2.94 (m, 1H), 2.70 - 2.54 (m, 2H), 2.45 - 2.16 (m, 2H), 1.48 (s, 9H). Resonances assigned to the O-alkylation byproduct are not reported.

*tert-butyl 2-(4-{2-[1-(5-chloro-1H-1,3-benzodiazol-2-yl)-5-hydroxy-3-[4-(trifluoromethyl)phenyl]-1H-pyrazol-4-yl]ethyl}phenoxy)acetate*. A mixture of ~3:1 mixture of methyl 4-{4-[2-(tert-butoxy)-2-oxoethoxy]phenyl}-2-[4-(trifluoromethyl)benzoyl]butanoate and methyl (2)-3-(2-{4-[2-(tert-butoxy)-2-oxoethoxy]phenyl}ethoxy)-3-[4-(trifluoromethyl)phenyl]prop-2-enoate (1.50 g, 3.1 mmol), 5-chloro-2-hydrazinyl-1H-1,3-benzodiazole (685 mg, 3.8 mmol), HOAc (3 mL) and EtOH (12 mL) was heated in the microwave at 130 °C for 3 h. Prep HPLC gave the title compound (260 mg, 13%).

$^1$H NMR (300MHz, $CD_3OD$) δ = 7.86-7.72 (s, 2H), 7.68 - 7.55 (m, 4H), 7.36 - 7.30 (m, 1H), 6.99 - 6.93 (m, 2H), 6.72 - 6.65 (m, 2H), 4.49 (s, 2H), 2.92 - 2.65 (m, 4H), 1.46 (s, 9H)

*2-(4-{2-[1-(5-chloro-1H-1,3-benzodiazol-2-yl)-5-hydroxy-3-[4-(trifluoromethyl)phenyl]-1H-pyrazol-4-yl]ethyl}phenoxy)acetic acid.* A solution of tert-butyl 2-(4-{2-[1-(5-chloro-1H-1,3-benzodiazol-2-yl)-5-hydroxy-3-[4-(trifluoromethyl)phenyl]-1H-pyrazol-4-yl]ethyl}phenoxy)acetate (260 mg, 0.42 mmol) in 1:1 $CH_2Cl_2$/TFA (6 mL) was stirred at rt for 1 h and concentrated. The residue was lyophilized from MeCN/5% aq HCl to give the title compound (220 mg, 94%).

LC-MS $t_R$ = 5.18 min, m/z 559, 557.

**2-[2-(4-{2-[1-(5-chloro-1H-1,3-benzodiazol-2-yl)-5-hydroxy-3-[4-(trifluoromethyl)phenyl]-1H-pyrazol-4-yl]ethyl}phenoxy)ethoxy]acetic acid**

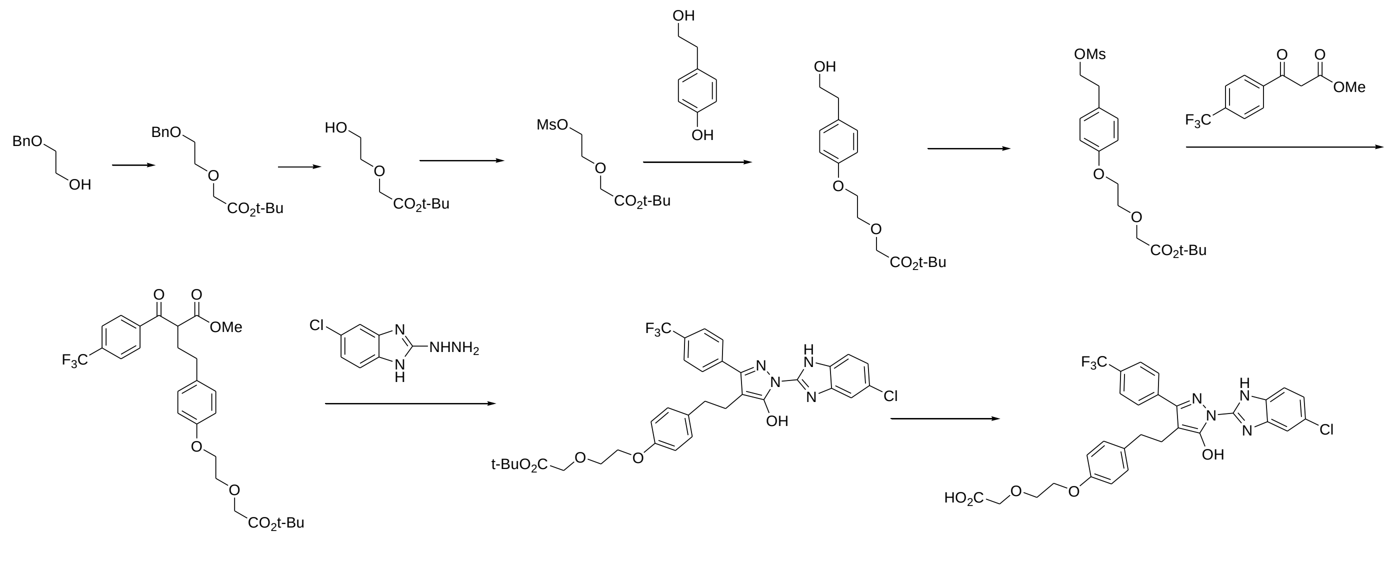

*tert-Butyl 2-[2-(benzyloxy)ethoxy]acetate.* To a stirred, ice-cold solution of 2-(benzyloxy)ethanol (3.19 g, 21.0 mmol) in dry DMF (20 mL) and dry THF (40 mL) was added 60% NaH in oil (1.26 g, 31.5 mmol) in several portions. The mixture was stirred in the ice bath for 15 min and t-butyl bromoacetate (6.2 mL, 41.9 mmol) was added over 2 min. The ice bath was allowed to melt, and the mixture was stirred at rt overnight. The mixture was recooled in an ice bath and brine (40 mL) was added dropwise over 5 min. The mixture was concentrated under reduced pressure to remove the bulk of the THF and the aqueous residue was extracted with EtOAc (2 x 50 mL). The combined EtOAc layer was washed with water (10 mL) and brine (10 mL), and dried over $Na_2SO_4$. Removal of the solvent left a mobile oil (8.81 g) which was purified by chromatography on an 80 g silica cartridge, eluted with a 0-50% EtOAc in hexanes gradient, to give the title compound (3.00 g, 54%).

$^1$H NMR (300MHz, $CDCl_3$) Shift = 7.42-7.30 (m, 5H), 4.60 (s, 2H), 4.12 (s, 2H), 3.85 - 3.64 (m, 4H), 1.60 - 1.32 (m, 9H)

*tert-Butyl 2-(2-hydroxyethoxy)acetate.* A solution of tert-butyl 2-[2-(benzyloxy)ethoxy]acetate (3.00 g, 11.3 mmol) in MeOH (100 mL) was shaken under $H_2$ (60 psi, Parr) in the presence of wet 10% Pd on C (1.00 g) for 4 h. The mixture was filtered through Celite, and the filtrate was concentrated to leave the title compound (2.06 g, quant.) as an oil.

$^1$H NMR (300MHz, $CDCl_3$) Shift = 4.01 (s, 2H), 3.78 - 3.60 (m, 4H), 3.11 - 2.97 (m, 1H), 1.47 (s, 9H).

*tert-butyl 2-[2-(methanesulfonyloxy)ethoxy]acetate*. To a solution of tert-butyl 2-(2-hydroxyethoxy)acetate (2.06 g, 11.7 mmol) and i-$Pr_2$NEt (4.2 mL, 23.4 mmol) in $CH_2Cl_2$ (40 mL) was added methanesulfonyl chloride (1.4 mL, 17.3 mmol). The ice bath was allowed to melt and the mixture was stirred at rt for 6 h. Additional i-$Pr_2$NEt (4.2 mL, 23.4 mmol) and methanesulfonyl chloride (1.4 mL, 17.3 mmol) were added and the mixture was stirred at rt for 18 h. The mixture was concentrated. The residue was taken up in EtOAc (80 mL) and 5% aq HCl (30 mL). The layers were separated and the EtOAc layer was washed with 5% aq HCl (30 mL) and brine (20 mL). The combined aqueous layer was back extracted with EtOAc (20 mL). The combined EtOAc layer was washed with brine (10 mL), dried over $Na_2SO_4$ and concentrated to leave the title compound (3.33 g, quant.) as a brown oil.

$^1$H NMR (300MHz, $CDCl_3$) Shift = 4.39 - 4.23 (m, 2H), 3.93 (s, 2H), 3.80 - 3.63 (m, 2H), 3.00 (s, 3H), 1.44 (s, 5H).

*tert-Butyl 2-{2-[4-(2-hydroxyethyl)phenoxy]ethoxy}acetate*. A mixture of 4-(2-hydroxyethyl)phenol (1.80 g, 13.0 mmol), $Cs_2CO_3$ (4.67 g, 14.3 mmol) and DMF (30 mL) was stirred at rt for 15 min and a solution of tert-butyl 2-[2-(methanesulfonyloxy)ethoxy]acetate (3.33 g, 13.0 mmol) in DMF (5 mL) was added. The mixture was stirred at rt for 1 d. Potassium iodide (2.17 g, 13.1 mmol) was added, and the mixture was stirred at 40 °C for 1 d. The mixture was diluted with EtOAc (175 mL) and washed with water (2 x 30 mL), 1 M aq NaOH (30 mL) and brine (30 mL). The water and aq NaOH washes were combined and back extracted with EtoAc. The combined EtOAc layer was dried over $Na_2SO_4$ and concentrated under reduced pressure to leave an oil (4.61 g). Chromatography on an 80 g silica cartridge, eluted with a 0-70% EtOAc in hexanes gradient, afforded the title compound (950 mg, 25%) as an oil.

$^1$H NMR (300MHz, $CDCl_3$) Shift = 7.20 - 7.04 (m, 2H), 6.93 - 6.77 (m, 2H), 4.18 - 4.11 (m, 2H), 4.09 (s, 2H), 3.95 - 3.86 (m, 2H), 3.85 - 3.76 (m, 2H), 2.83 - 2.74 (m, 2H), 1.47 (s, 9H).

*tert-Butyl 2-(2-{4-[2-(methanesulfonyloxy)ethyl]phenoxy}ethoxy)acetate*. To a solution of tert-butyl 2-{2-[4-(2-hydroxyethyl)phenoxy]ethoxy}acetate (950 mg, 3.2 mmol) and i-$Pr_2$NEt (1.8 mL, 9.7 mmol) in $CH_2Cl_2$ (30 mL) was added methanesulfonyl chloride (0.60 mL, 7.7 mmol). The ice bath was allowed to melt, and the mixture was stirred at rt for 6 h and concentrated. The residue was taken up in EtOAc (90 mL), washed with 5% aq HCl (2 x 20 mL) and brine (20 mL), and dried over $Na_2SO_4$. Removal of the solvent left the title compound (1.14 g, 95%) as a brown oil.

$^1$H NMR (300MHz, $CDCl_3$) Shift = 7.18 - 7.08 (m, 2H), 6.92 - 6.81 (m, 2H), 4.37 (t, J=6.7 Hz, 2H), 4.20 - 4.10 (m, 2H), 4.05 (s, 2H), 3.97 - 3.85 (m, 2H), 2.98 (t, J=7.0 Hz, 2H), 2.83 (s, 3H), 1.48 (s, 9H).

*Methyl 4-(4-{2-[2-(tert-butoxy)-2-oxoethoxy]ethoxy}phenyl)-2-[4-(trifluoromethyl)benzoyl] butanoate.* A mixture of methyl 3-oxo-3-[4-(trifluoromethyl)phenyl]propanoate (750 mg, 3.0 mmol), tert-butyl 2-(2-{4-[2-(methanesulfonyloxy)ethyl]phenoxy}ethoxy)acetate (1.14 g, 3.0 mmol), powdered $K_2CO_3$ (420 mg, 3.0 mmol), KI (500 mg, 3.0 mmol) and dry DMF (12 mL) was stirred in a 70 °C oil bath under $N_2$ for 8 h. The mixture was diluted with EtOAc (90 mL), washed with 5% aq HCl (2 x 20 mL) and brine (20 mL), and dried over $Na_2SO_4$. Removal of the solvent left a mobile oil (1.93 g) which was purified by chromatography on an 80 g silica cartridge, eluted with a 0-30% EtOAc in hexanes gradient, to give a 2:1 mixture of the title compound and the O-alkylation product methyl (2Z)-3-[2-(4-{2-[2-(tert-butoxy)-2-oxoethoxy]ethoxy}phenyl)ethoxy]-3-[4-(trifluoromethyl)phenyl]prop-2-enoate (730 mg, 45%) as an oil.

$^1H$ NMR (300MHz, $CDCl_3$) Shift = 8.00 - 7.88 (m, 2H), 7.79 - 7.64 (m, 2H), 7.04 (d, J=8.8 Hz, 2H), 6.84 (d, J=8.3 Hz, 2H), 4.22 - 4.11 (m, 2H), 4.08 (s, 2H), 3.94 - 3.87 (m, 2H), 3.68 (s, 3H), 3.07 - 2.95 (m, 1H), 2.67 - 2.54 (m, 2H), 2.39 - 2.18 (m, 2H), 1.46 (s, 9H). Resonances assigned to the O-alkylation byproduct are not reported.

*tert-Butyl 2-[2-(4-{2-[1-(5-chloro-1H-1,3-benzodiazol-2-yl)-5-hydroxy-3-[4-(trifluoromethyl) phenyl]-1H-pyrazol-4-yl]ethyl} phenoxy)ethoxy]acetate.* A mixture of methyl 4-(4-{2-[2-(tert-butoxy)-2-oxoethoxy]ethoxy}phenyl)-2-[4-(trifluoromethyl)benzoyl]butanoate (730 mg, 1.4 mmol), 5-chloro-2-hydrazinyl-1H-1,3-benzodiazole (178 mg, 0.97 mmol), HOAc (1 mL) and MeOH (3 mL) was heated in the microwave at 130 °C for 3 h. Prep HPLC afforded the title compound (132 mg, 14%) as a solid.

$^1H$ NMR (300MHz, $CD_3OD$) Shift = 7.82 - 7.55 (m, 6H), 7.45 - 7.34 (m, 1H), 6.98 - 6.88 (m, 2H), 6.75 - 6.65 (m, 2H), 4.08 (s, 2H), 4.07 - 3.99 (m, 2H), 3.87 - 3.81 (m, 2H), 2.92 - 2.83 (m, 2H), 2.78 - 2.68 (m, 2H), 1.46 (s, 9H)

LC-MS $t_R$ 6.33 min, m/z 657, 601.

*2-[2-(4-{2-[1-(5-Chloro-1H-1,3-benzodiazol-2-yl)-5-hydroxy-3-[4-(trifluoromethyl)phenyl]-1H-pyrazol-4-yl]ethyl}phenoxy) ethoxy]acetic acid.* To a stirred solution of tert-butyl 2-[2-(4-{2-[1-(5-chloro-1H-1,3-benzodiazol-2-yl)-5-hydroxy-3-[4-(trifluoromethyl)phenyl]-1H-pyrazol-4-yl]ethyl}phenoxy)ethoxy]acetate (128 mg, 0.19 mmol) in $CH_2Cl_2$ (3 mL) was added $CF_3CO_2H$ (3 mL). The mixture was stirred at rt for 0.5 h and concentrated. The residue was lyophilized from MeCN/5% aq HCl to give the HCl salt of the title compound (114 mg, 92%) as a tan solid.

$^1H$ NMR (300MHz, DMSO-$d_6$) Shift = 7.91 - 7.73 (m, 4H), 7.63 - 7.52 (m, 2H), 7.26 (dd, J=2.2, 8.8 Hz, 1H), 7.03 (d, J=8.8 Hz, 2H), 6.77 (d, J=8.3 Hz, 2H), 4.07 (s, 2H), 4.04 - 3.98 (m, 2H), 3.79 - 3.70 (m, 2H), 2.84 - 2.64 (m, 4H).

LC-MS $t_R$ 5.28 min, m/z 623, 601.

**1-(5-chloro-1H-1,3-benzodiazol-2-yl)-4-[2-(4-fluorophenyl)ethyl]-3-(piperidin-4-yl)-1H-pyrazol-5-ol**

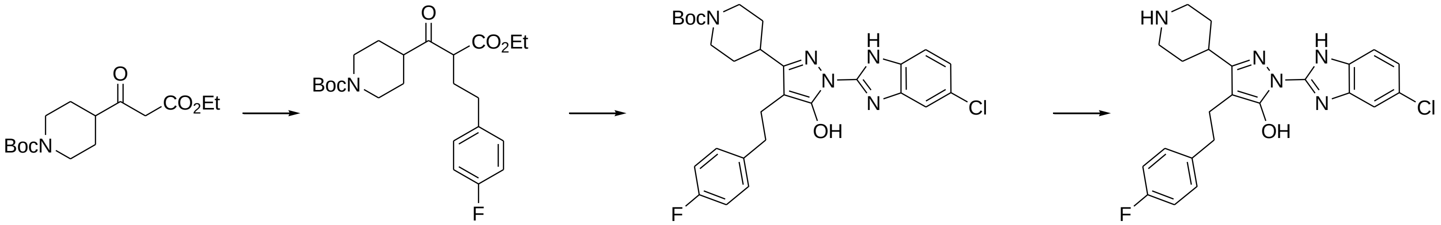


*tert-butyl 4-{3-ethoxy-2-[2-(4-fluorophenyl)ethyl]-3-oxopropanoyl}piperidine-1-carboxylate*. A mixture of tert-butyl 4-(3-ethoxy-3-oxopropanoyl)piperidine-1-carboxylate (3.72 g, 12.4 mmol), 1-(2-bromoethyl)-4-fluorobenzene (2.91 g, 14.3 mmol), $K_2CO_3$ (1.89 g, 13.7 mmol), KI (2.27 g, 13.7 mmol) and DMF (30 mL) was stirred at 70 $^oC$ for 8 h. The mixture was diluted with EtOAc (175 mL), washed with 5% aq HCl (30 mL), water (2 x 30 mL) and brine (30 mL), and dried over $Na_2SO_4$. Removal of the solvent left a brown oil (6.85 g). Chromatography on an 80 g silica cartridge, eluted with a 0-40% EtOAc in hexanes gradient, gave the title compound (3.59 g, 68%) as a pale yellow oil.

$^1H$ NMR ($CDCl_3$) 7.20-7.03 (m, 2H), 7.01-6.85 (m, 2H), 4.25-3.93 (m, 6H), 3.60-3.52 (m, 1H), 2.80-2.45 (m, 6H), 2.20-2.05 (m, 1H), 1.88-1.45 (m, 2H), 1.40 (s, 9H), 1.18-1.27 (m, 3H). LC-MS $t_R$ = 5.73 min, m/z 322 [M + Na+].

*tert-butyl 4-[1-(5-chloro-1H-1,3-benzodiazol-2-yl)-4-[2-(4-fluorophenyl)ethyl]-5-hydroxy-1H-pyrazol-3-yl]piperidine-1-carboxylate.* A mixture of tert-butyl 4-{3-ethoxy-2-[2-(4-fluorophenyl)ethyl]-3-oxopropanoyl}piperidine-1-carboxylate (1.70 g, 4.0 mmol), 5-chloro-2-hydrazinyl-1H-1,3-benzodiazole (920 mg, 5.0 mmol), HOAc (3 mL) and EtOH (12 mL) was heated in the microwave at 130 $^oC$ for 3 h. Prep HPLC gave the title compound (1.02 g, 58%) as solid.

$^1H$ NMR ($CD_3OD$) δ: 7.52-7.63 (m, 2H), 7.31 (dd, J=8.6, 2.0 Hz, 1H), 7.14-7.23 (m, 2H), 6.93-7.05 (m, 2H), 4.10 (br d, J=13.3 Hz, 2H), 2.62-2.90 (m, 6H), 2.42-2.58 (m, 1H), 1.70-1.48 (br dd, J=17.5, 4.3 Hz, 4H), 1.46 (s, 9H).

$^{19}F$ NMR ($CD_3OD$) δ: -77.45, -119.47.

LC-MS $t_R$ 5.72 min, m/z 542, 540, 442, 440.

HRMS calc'd for $C_{28}H_{32}ClFN_5O_3$ 540.2172, found 540.217

*1-(5-chloro-1H-1,3-benzodiazol-2-yl)-4-[2-(4-fluorophenyl)ethyl]-3-(piperidin-4-yl)-1H-pyrazol-5-ol.* A solution of tert-butyl 4-[1-(5-chloro-1H-1,3-benzodiazol-2-yl)-4-[2-(4-fluorophenyl)ethyl]-5-hydroxy-1H-pyrazol-3-yl]piperidine-1-carboxylate (1.02 g, 1.90 mmol) in 3:1 $CH_2Cl_2$/TFA (8 mL) was stirred at rt for 1.5 h and concentrated. The residue was lyophilized from MeCN/5% aq HCl to give the HCl salt of the title compound (829 mg, quant) as a grey solid.

LC-MS $t_R$ = 3.90 min, m/z 442, 440.

**2-{2-[2-(3-{4-[1-(5-chloro-1H-1,3-benzodiazol-2-yl)-4-[2-(4-fluorophenyl)ethyl]-5-hydroxy-1H-pyrazol-3-yl]phenyl} propoxy) ethoxy]ethoxy}acetic acid**

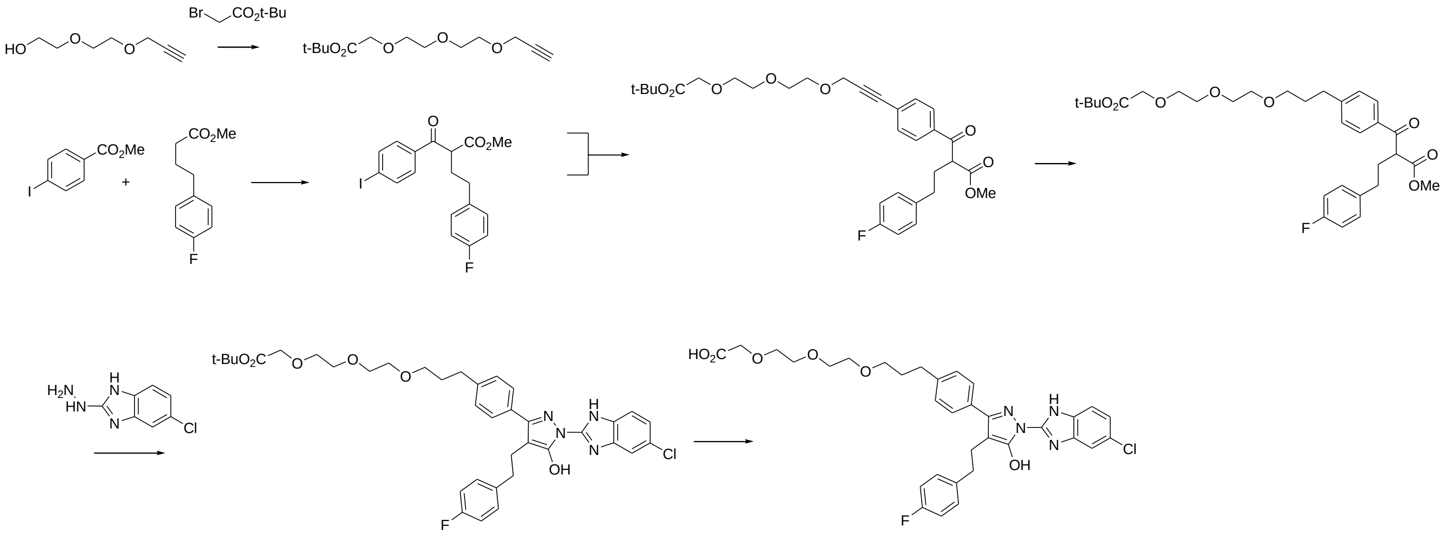


*tert-butyl 2-{2-[2-(prop-2-yn-1-yloxy)ethoxy]ethoxy}acetate*. To a stirred, ice-cold suspension of 60% NaH in oil (700 mg, 17.3 mmol) in dry THF (20 mL) was added dropwise over 5 min a solution of 2-[2-(prop-2-yn-1-yloxy)ethoxy]ethan-1-ol (1.78 g, 12.3 mmol) in dry THF (10 mL). The mixture was stirred at rt for 1 h, recooled in an ice bath and treated with t-butyl bromoacetate (3.65 mL, 24.7 mmol) dropwise over 3 min. The mixture was allowed to warm to rt and stirred for 18 h. The mixture was poured into ice-cold 5% aq HCl (50 mL) and extracted with EtOAc (2 x 40 mL). The combined organic layer was washed with brine (15 mL), dried over $Na_2SO_4$ and concentrated to leave a yellow oil (5.30 g). Chromatography on silica gel, eluted with an ethyl acetate hexane gradient, gave the title compound (1.08 g, 86%) as a colorless oil.

$^1$H NMR (300MHz, $CDCl_3$) δ = 4.18 (s, 2H), 3.96 (d, *J*=0.9 Hz, 1H), 3.65 (br. s., 8H), 2.44 - 2.32 (m, 1H), 1.42 (s, 9H).

*Methyl 4-(4-fluorophenyl)-2-(4-iodobenzoyl)butanoate*. An oven-dried flask, equipped with a stir bar was charged with methyl 4-iodobenzoate (4.77 g, 18.2 mmol), methyl 4-(4-fluorophenyl)butanoate (2.98 g, 15.2 mmol) and 60% NaH in oil (1.30 g, 31.9 mmol). The flask was sealed with a septum and dry THF (30 mL) was introduced by syringe., followed by MeOH (2 drops). The mixture was stirred at 70 °C under $N_2$ for 8 h, cooled to rt and poured into ice-cold 2.5% aq HCl. The mixture was extracted with EtOAc (3 x 35 mL). The combined EtOAc layer was washed with brine (15 mL), dried over $Na_2SO_4$ and concentrated to leave a wet solid (7.85 g). Chromatography on an 80 g silica cartridge, eluted with a 0-25% EtOAc in hexanes gradient, afforded the title compound (3.77 g, 58%) as a colorless oil.

$^1$H NMR (300MHz, $CDCl_3$) δ = 7.88 - 7.75 (m, 2H), 7.63 - 7.54 (m, 2H), 7.18 - 7.07 (m, 2H), 6.96 (s, 2H), 4.21 (t, *J*=7.0 Hz, 1H), 3.68 (s, 3H), 2.70 - 2.54 (m, 2H), 2.41 - 2.16 (m, 2H).

*Methyl 2-{4-[3-(2-{2-[2-(tert-butoxy)-2-oxoethoxy]ethoxy}ethoxy)prop-1-yn-1-yl]benzoyl}-4-(4-fluorophenyl)butanoate*. A flask was charged with methyl 4-(4-fluorophenyl)-2-(4-iodobenzoyl)butanoate (900 mg, 2.1 mmol), tert-butyl 2-{2-[2-(prop-2-yn-1-yloxy)ethoxy]ethoxy}acetate (819 mg, 2.2 mmol), CuI (41 mg, 0.21 mmol), $Pd(PPh_3)_2Cl_2$ (150 mg, 0.21 mmol) and sealed with a septum. The flask was evacuated/refilled with $N_2$ (3x) and dry $CH_2Cl_2$ (10 mL) and $Et_3N$ (2.5 mL) were added via syringe. The flask was evacuated/refilled with $N_2$ (3x), stirred at rt for 2 d and concentrated. The residue was taken up in EtOAc (100 mL), washed with 5% aq HCl (2 x 10 mL) and brine (10 mL), and dried over $Na_2SO_4$. Removal of the solvent left a brown oil (1.84 g) which was chromatographed on a 40 g silica cartridge, eluted with a 0-60% EtOAc in hexanes gradient, to provide the title compound (760 mg, 64%) as an oil.

$^1$H NMR (300MHz, $CDCl_3$) δ = 7.86-7.78 (m, 2H), 7.55 - 7.46 (m, 2H), 7.15 - 7.05 (m, 2H), 7.03 - 6.87 (m, 2H), 4.45 (s, 2H), 4.31 - 4.20 (m, 1H), 4.03 (s, 2H), 3.80-3.64 (m, 11H), 2.74 - 2.57 (m, 2H), 2.42 - 2.17 (m, 2H), 1.42 (s, 9H).

*Methyl 2-{4-[3-(2-{2-[2-(tert-butoxy)-2-oxoethoxy]ethoxy}ethoxy)propyl]benzoyl}-4-(4-fluorophenyl)butanoate*. A solution of methyl 2-{4-[3-(2-{2-[2-(tert-butoxy)-2-oxoethoxy]ethoxy}ethoxy)prop-1-yn-1-yl]benzoyl}-4-(4-fluorophenyl)butanoate (760 mg, 1.4 mmol) in MeOH (20 mL) was stirred with 10% Pd on C (cat. qty.) under $H_2$ (1 atm, balloon) at rt for 4 h. The flask was flushed with $N_2$ and the mixture was filtered through Celite. The filtrate was concentrated to leave a brown oil (730 mg) which was purified by chromatography on a 24 g silica cartridge, eluted with a 0-70% EtOAc in hexanes gradient, to give the title compound (460 mg, 60%) as a brown oil.

$^1$H NMR (300MHz, $CDCl_3$) δ = 7.86 - 7.76 (m, 2H), 7.35-7.24 (m, 2H), 7.18 - 7.07 (m, 2H), 7.01 - 6.89 (m, 2H), 4.33 - 4.20 (m, 1H), 4.02 (s, 2H), 3.77-3.62 (m, 9H), 3.62 - 3.55 (m, 2H), 3.51 - 3.38 (m, 2H), 2.80 - 2.69 (m, 2H), 2.67 - 2.58 (m, 2H), 2.37 - 2.21 (m, 2H), 1.98 - 1.83 (m, 2H), 1.46 (s, 9H)

*tert-butyl 2-{2-[2-(3-{4-[1-(5-chloro-1H-1,3-benzodiazol-2-yl)-4-[2-(4-fluorophenyl)ethyl]-5-hydroxy-1H-pyrazol-3-yl]phenyl}propoxy)ethoxy]ethoxy}acetate*. A mixture of methyl 2-{4-[3-(2-{2-[2-(tert-butoxy)-2-oxoethoxy]ethoxy}ethoxy)propyl]benzoyl}-4-(4-fluorophenyl)butanoate (460 mg, 0.82 mmol), 5-chloro-2-hydrazinyl-1H-1,3-benzodiazole (188 mg, 1.03 mmol), HOAc (0.5 mL) and MeOH (1.5 mL) was heated in the microwave at 130 °C for 3 h. Prep HPLC gave the title compound (190 mg, 33%) as a solid.

$^1$H NMR (300MHz, $CD_3OD$) δ = 7.62 - 7.47 (m, 2H), 7.47 - 7.39 (m, 2H), 7.34 - 7.22 (m, 3H), 7.10 - 6.98 (m, 2H), 6.91 - 6.78 (m, 2H), 4.0 (s, 2H), 3.71 - 3.61 (m, 6H), 3.60 - 3.53 (m, 2H), 3.49 - 3.41 (m, 2H), 2.81 - 2.64 (m, 6H), 1.95 - 1.77 (m, 2H), 1.44 (s, 9H). LC-MS $t_R$ = 6.15 min, m/z 693, 637.

*2-{2-[2-(3-{4-[1-(5-chloro-1H-1,3-benzodiazol-2-yl)-4-[2-(4-fluorophenyl)ethyl]-5-hydroxy-1H-pyrazol-3-yl]phenyl}propoxy)ethoxy]ethoxy}acetic acid*. A solution of tert-butyl 2-{2-[2-(3-{4-[1-(5-chloro-1H-1,3-benzodiazol-2-yl)-4-[2-(4-fluorophenyl)ethyl]-5-hydroxy-1H-pyrazol-3-yl]phenyl}propoxy)ethoxy]ethoxy}acetate (165 mg, 0.24 mmol) in 1:1 $CH_2Cl_2$/TFA (6 mL) was stirred at rt for 3 h and concentrated. The residue was lyophilized from MeCN/5% aq HCl to give (136 mg, 90%) as a tan solid.

$^1$H NMR (300MHz, $CD_3OD$) δ = 7.64 - 7.53 (m, 3H), 7.50 - 7.43 (m, 1H), 7.37 - 7.29 (m, 3H), 7.13 - 7.03 (m, 2H), 6.94 - 6.83 (m, 2H), 4.14 (s, 2H), 3.78 - 3.56 (m, 8H), 3.55 - 3.45 (m, 2H), 2.93 - 2.67 (m, 6H), 2.02 - 1.82 (m, 2H)

**3-[4-({2-[2-(2-aminoethoxy)ethoxy]ethoxy}methyl)phenyl]-1-(5-chloro-1H-1,3-benzodiazol-2-yl)-4-[2-(4-fluorophenyl)ethyl]-1H-pyrazol-5-ol**

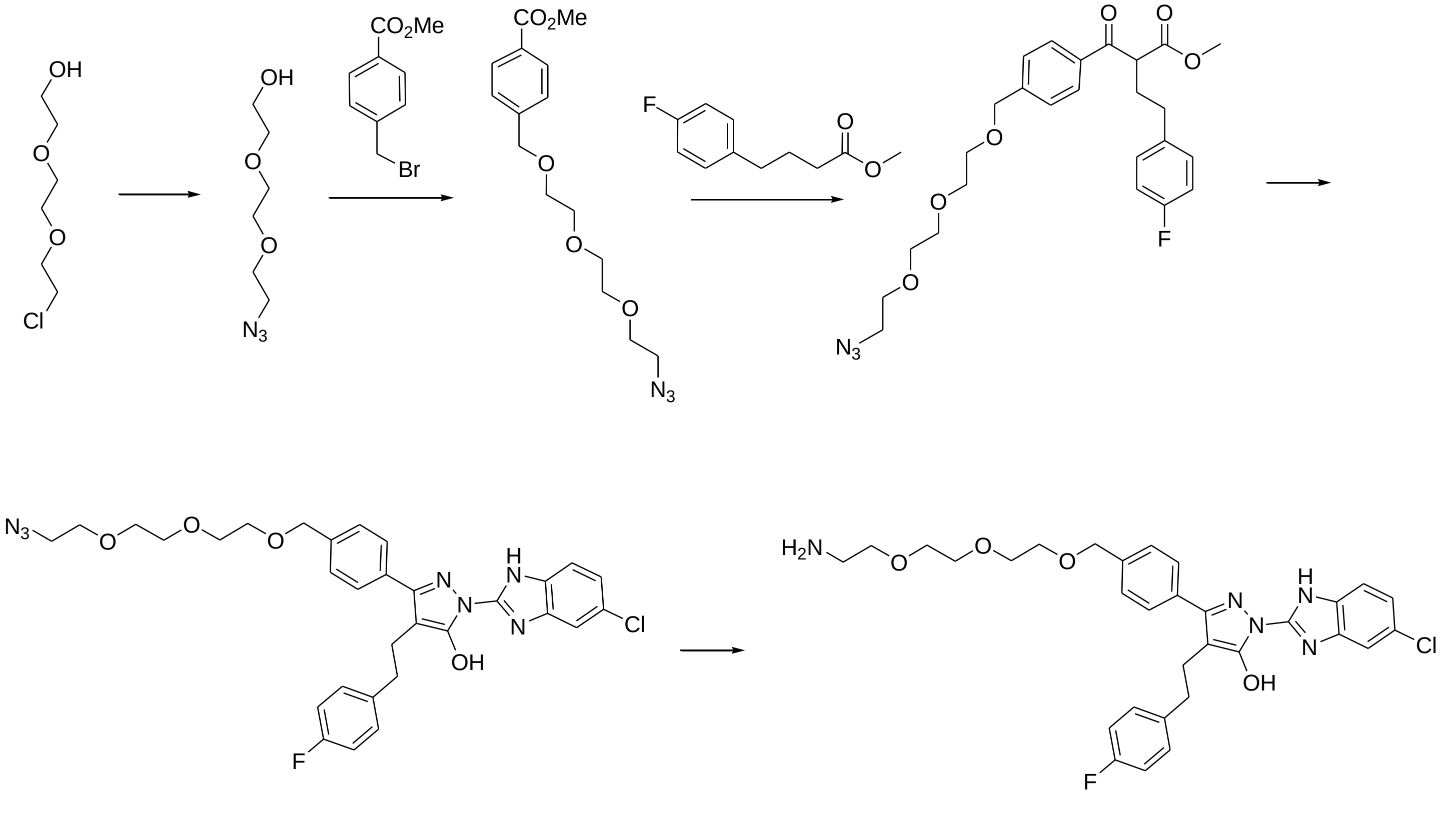

*2-{2-[2-azidoethoxy]ethoxy}ethan-1-ol*. A mixture of 2-[2-(2-chloroethoxy)ethoxy]ethan-1-ol (3.75 g, 22.2 mmol), $NaN_3$ (3.60 g, 55.5 mmol) and water (15 mL) was stirred at 80 °C for 1 d, cooled to rt, diluted with 1 M aq NaOH (5 mL, 5.0 mmol) and extracted with ether (3 x 35 mL). The combined ether layer was dried over $Na_2SO_4$ and concentrated to leave the title compound (2.50 g, 64%) as a colorless oil.

$^1$H NMR (300MHz, $CDCl_3$) δ = 3.75 - 3.63 (m, 8H), 3.62-3.56 (m, 2H), 3.42-3.35 (m, 2H).

*Methyl 4-({2-[2-(2-azidoethoxy)ethoxy]ethoxy}methyl)benzoate*. To a stirred solution of methyl 4-(bromomethyl)benzoate (1.57 g, 6.9 mmol) and 2-{2-[2-azidoethoxy]ethoxy}ethan-1-ol (1.26 g, 7.2 mmol) in dry THF (10 mL) was added 60% NaH in oil (330 mg, 8.3 mmol). The mixture was stirred at rt for 5 d, diluted with EtOAc (90 mL), washed with 5% aq HCl (20 mL), satd aq $NaHCO_3$ (20 mL) and brine (20 mL), and dried over $Na_2SO_4$. Removal of the solvent left an oil (2.40 g) which was chromatographed on a 40 g silica cartridge, eluted with a 0-80% EtOAc in hexanes gradient, to give the title compound (810 mg, 36%) as a colorless oil.

$^1$H NMR (300MHz, $CDCl_3$) δ = 8.01 (d, *J*=7.9 Hz, 2H), 7.41 (d, *J*=7.9 Hz, 2H), 4.62 (s, 2H), 3.72-3.60 (m, 10H), 3.43 - 3.31 (m, 2H).

*Methyl 2-[4-({2-[2-(2-azidoethoxy)ethoxy]ethoxy}methyl)benzoyl]-4-(4-fluorophenyl)butanoate*. To a mixture of methyl 4-(4-fluorophenyl)butanoate (280 mg, 1.4 mmol) and methyl 4-({2-[2-(2-azidoethoxy)ethoxy]ethoxy}methyl)benzoate (810 mg, 2.5 mmol) was added 60% NaH in oil (285 mg, 7.1 mmol), followed by dry THF (5 mL) and MeOH (1 drop). The mixture was heated at reflux under $N_2$ for 5 h, diluted with EtOAc (90 mL), washed with 5% aq HCl (15 mL) and brine (15 mL), and dried over $Na_2SO_4$. Removal of the solvent left an oil (1.08 g) which was purified by chromatography on a 40 g silica cartridge, eluted with a 0-100% EtOAc in hexanes gradient, to provide the title compound (170 mg, 24%).

$^1$H NMR (300MHz, $CDCl_3$) δ = 7.92 - 7.80 (m, 2H), 7.48 - 7.35 (m, 2H), 7.11 (dd, *J*=5.5, 8.6 Hz, 2H), 7.03 - 6.85 (m, 2H), 4.63 (s, 2H), 4.34 - 4.21 (m, 1H), 3.78 - 3.61 (m, 13H), 3.47 - 3.29 (m, 2H), 2.68-2.60 (m, 2H), 2.40 - 2.19 (m, 2H).

*3-[4-({2-[2-(2-azidoethoxy)ethoxy]ethoxy}methyl)phenyl]-1-(5-chloro-1H-1,3-benzodiazol-2-yl)-4-[2-(4-fluorophenyl)ethyl] 1H-pyrazol-5-ol*. A mixture of methyl 2-[4-({2-[2-(2-azidoethoxy)ethoxy]ethoxy}methyl)benzoyl]-4-(4-fluorophenyl)butanoate (170 mg, 0.35 mmol), 5-chloro-2-hydrazinyl-1H-1,3-benzodiazole (67 mg, 0.37 mmol), $TsOH.H_2O$ (14 mg, 0.07 mmol) and MeOH (4 mL) was stirred at 70 °C under $N_2$ for 3 d. Prep HPLC gave the title compound (94 mg, 43%) as a tan solid.

$^1$H NMR (300MHz, $CD_3OD$) δ = 7.64 - 7.35 (m, 6H), 7.32 - 7.19 (m, 1H), 7.05 (dd, *J*=5.5, 8.6 Hz, 2H), 6.94 - 6.79 (m, 2H), 4.58 (s, 2H), 3.77 - 3.56 (m, 10H), 3.43 - 3.20 (m, 2H), 2.78 (s, 4H).

LC-MS $t_R$ 5.43 min, m/z 622, 620.

*3-[4-({2-[2-(2-aminoethoxy)ethoxy]ethoxy}methyl)phenyl]-1-(5-chloro-1H-1,3-benzodiazol-2-yl)-4-[2-(4-fluorophenyl)ethyl]-1H-pyrazol-5-ol.* To a stirred solution of 3-[4-({2-[2-(2-azidoethoxy)ethoxy]ethoxy}methyl)phenyl]-1-(5-chloro-1H-1,3-benzodiazol-2-yl)-4-[2-(4-fluorophenyl)ethyl]-1H-pyrazol-5-ol (86 mg, 0.14 mmol) in dry THF (3 mL) was added 1 M $Me_3P$ in THF (0.42 mL, 0.42 mmol). The mixture was stirred at rt under $N_2$ for 2 h and water (0.3 mL) was added. The mixture was stirred at rt for 2 d and 1M aq NaOH (0.5, 0.5 mmol) was added. The mixture was stirred at rt for 3 h, diluted with HOAc (1 mL) and purified by prep HPLC to give the TFA salt of the title compound (76 mg, 77%) as a white solid.

$^1$H NMR (300MHz, $CD_3OD$) δ = 7.65 - 7.38 (m, 6H), 7.35 - 7.23 (m, 1H), 7.14 - 7.00 (m, 2H), 6.95 - 6.77 (m, 2H), 4.61 (s, 2H), 3.80 - 3.59 (m, 12H), 3.18 - 3.06 (m, 2H), 2.81 (s, 4H).

LC-MS $t_R$ = 4.23 min, m/z 594.

### Synthesis of PROTACs

**N-(2-{2-[2-({4-[1-(5-chloro-1H-1,3-benzodiazol-2-yl)-4-[2-(4-fluorophenyl)ethyl]-5-hydroxy-1H-pyrazol-3-yl]phenyl}methoxy)ethoxy]ethoxy}ethyl)-2-{[2-(2,6-dioxopiperidin-3-yl)-1-oxo-2,3-dihydro-1H-isoindol-4-yl]oxy}acetamide (FC-12988)**

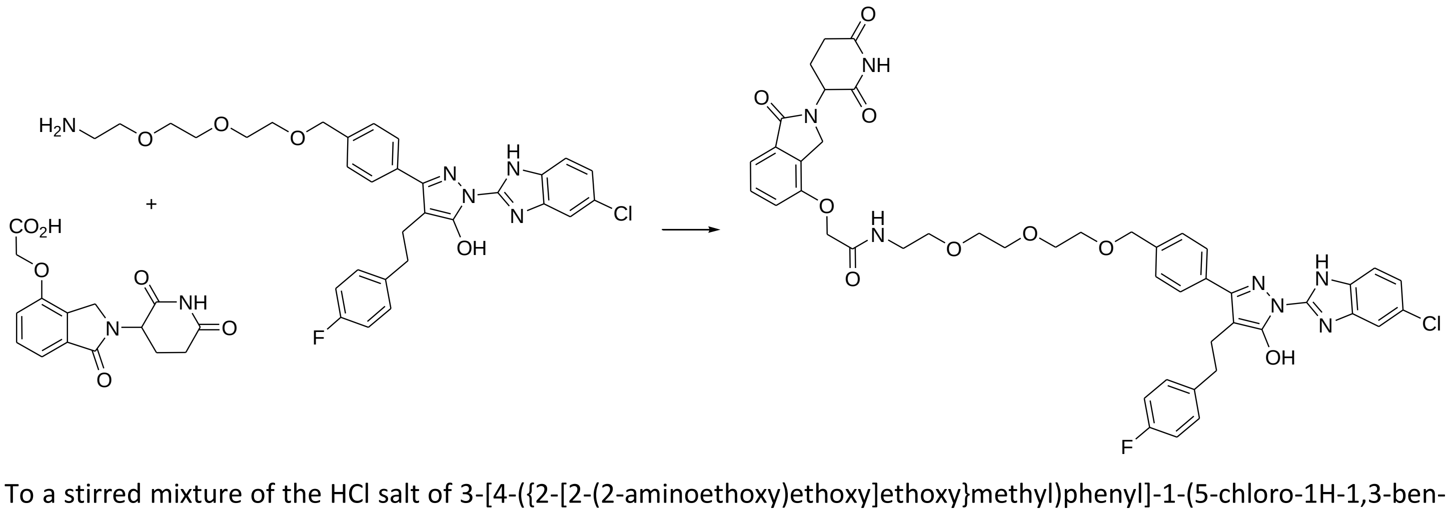

To a stirred mixture of the HCl salt of 3-[4-({2-[2-(2-aminoethoxy)ethoxy]ethoxy}methyl)phenyl]-1-(5-chloro-1H-1,3-benzodiazol-2-yl)-4-[2-(4-fluorophenyl)ethyl]-1H-pyrazol-5-ol (55 mg, 82 µmol), 2-{[2-(2,6-dioxopiperidin-3-yl)-1-oxo-2,3-dihydro-1H-isoindol-4-yl]oxy}acetic acid (53 mg, 0.17 mmol), i-$Pr_2NEt$ (80 µL, 0.45 mmol), dry $CH_2Cl_2$ (1 mL) and dry DMF (1 mL) was added solid HATU (63 mg, 0.17 mmol). The mixture was stirred at rt under $N_2$ for 2 h and concentrated under reduced pressure to remove $CH_2Cl_2$. The residue was purified by prep HPLC, followed by lyophilization from MeCN/5% aq HCl, to give the HCl salt of the title compound (33 mg, 43%) as an off-white solid.

$^1$H NMR (300MHz, $CD_3OD$) Shift = 7.80 - 7.66 (m, 2H), 7.63 - 7.30 (m, 7H), 7.15 - 6.99 (m, 3H), 6.89 (s, 2H), 5.12 (dd, J=5.1, 13.4 Hz, 1H), 4.63 - 4.61 (m, 2H), 4.59 - 4.57 (m, 2H), 4.47 - 4.43 (m, 2H), 3.69 - 3.66 (m, 4H), 3.63 - 3.56 (m, 6H), 3.50 - 3.43 (m, 2H), 2.96 - 2.82 (m, 3H), 2.80 - 2.65 (m, 3H), 2.51 - 2.29 (m, 1H), 2.19 - 1.99 (m, 1H).

$^{19}$F NMR (282MHz, $CD_3OD$) Shift = -77.79, -119.84.

LC-MS $t_R$ = 4.33 min, m/z 894.

HRMS Calc'd for $C_{46}H_{46}ClFN_7O_9$: 894.302408; found: 894.304560.

**2-[2-(4-{2-[1-(5-chloro-1H-1,3-benzodiazol-2-yl)-5-hydroxy-3-[4-(trifluoromethyl)phenyl]-1H-pyrazol-4-yl]ethyl}phenoxy)ethoxy]-N-(2-{[2-(2,6-dioxopiperidin-3-yl)-1,3-dioxo-2,3-dihydro-1H-isoindol-4-yl]amino}ethyl)acetamide (FC-13182)**

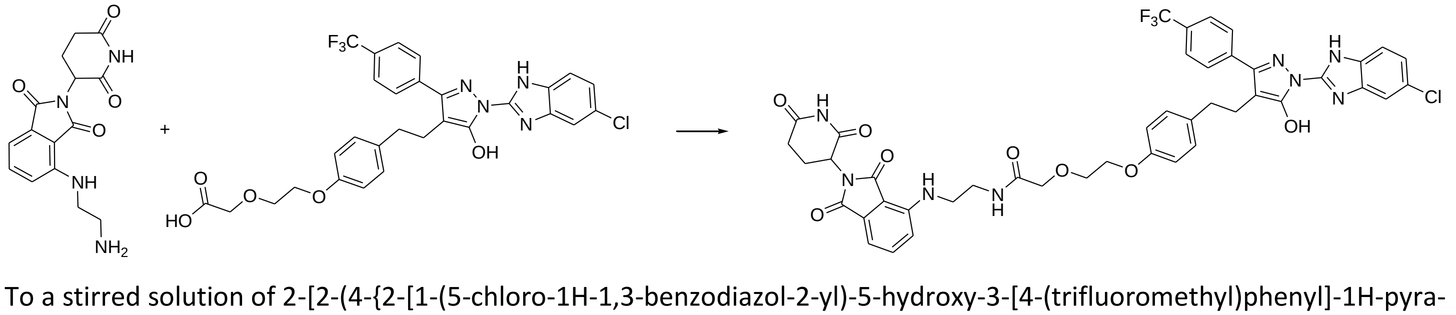

To a stirred solution of 2-[2-(4-{2-[1-(5-chloro-1H-1,3-benzodiazol-2-yl)-5-hydroxy-3-[4-(trifluoromethyl)phenyl]-1H-pyrazol-4-yl]ethyl}phenoxy)ethoxy]acetic acid HCl salt (26 mg, 41 µmol), 4-[(2-aminoethyl)amino]-2-(2,6-dioxopiperidin-3-yl)-2,3-dihydro-1H-isoindole-1,3-dione (14.5 mg, 41 µmol), HOBt.$H_2O$ (7 mg, 45 µmol) and i-$Pr_2NEt$ (40 µL, 0.22 mmol) in dry DMF (1 mL) was added EDC.HCl (16 mg, 84 µmol). The mixture was stirred overnight at rt and purified by prep HPLC to give the TFA salt of the title compound (23 mg, 50%) as a yellow solid.

$^1H$ NMR (300MHz, DMSO-$d_6$) Shift = 8.03 - 7.95 (m, 1H), 7.89 - 7.76 (m, 4H), 7.62 - 7.48 (m, 3H), 7.25 (d, J=2.2 Hz, 1H), 7.16 (d, J=8.8 Hz, 1H), 7.06 - 6.96 (m, 3H), 6.82-6.63 (m, 3H), 5.07 - 4.96 (m, 1H), 4.04 (br s, 2H), 3.92 (s, 2H), 3.78 - 3.69 (m, 2H), 3.42 - 3.22 (m, 4H), 2.95 - 2.77 (m, 1H), 2.76-2.63 (m, 4H), 2.61 - 2.54 (m, 1H), 2.44 - 2.39 (m, 1H), 2.04 - 1.91 (m, 1H).

$^{19}F$ NMR (282MHz, DMSO-$d_6$) Shift = -61.09, -74.88.

LC-MS $t_R$ 5.38 min, m/z 901, 899.

HRMS Calc'd for $C_{44}H_{39}ClF_3N_8O_8$: 899.2526; found: 899.2529.

**2-{2-[2-(3-{4-[1-(5-chloro-1H-1,3-benzodiazol-2-yl)-4-[2-(4-fluorophenyl)ethyl]-5-hydroxy-1H-pyrazol-3-yl]phenyl}propoxy)ethoxy]ethoxy}-N-(2-{[2-(2,6-dioxopiperidin-3-yl)-1,3-dioxo-2,3-dihydro-1H-isoindol-4-yl]amino}ethyl)acetamide (FC-13818)**

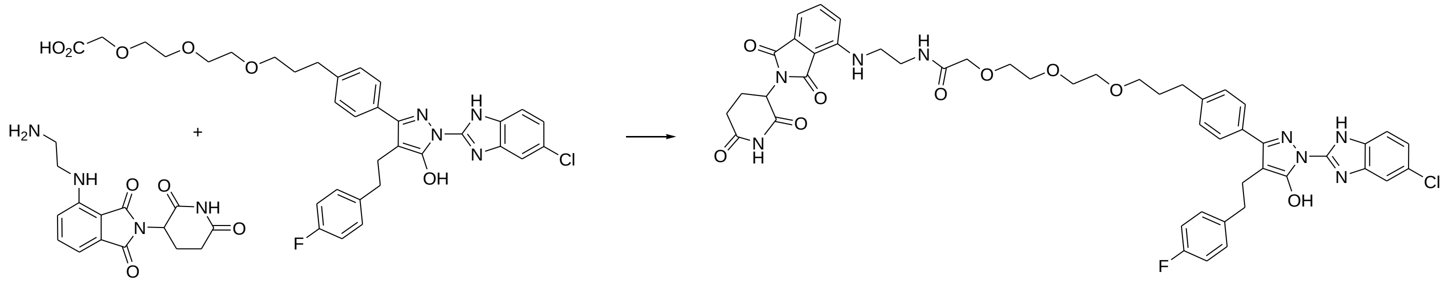

To a stirred solution of the TFA salt of 2-{2-[2-(3-{4-[1-(5-chloro-1H-1,3-benzodiazol-2-yl)-4-[2-(4-fluorophenyl)ethyl]-5-hydroxy-1H-pyrazol-3-yl]phenyl}propoxy)ethoxy]ethoxy}acetic acid (30 mg, 47 µmol), the HCl salt of 4-[(2-aminoethyl)amino]-2-(2,6-dioxopiperidin-3-yl)-2,3-dihydro-1H-isoindole-1,3-dione (16.5 mg, 46 µmol), HOBt.$H_2O$ (7.5 mg, 49 mol) and i-$Pr_2NEt$ (45 µL, 0.25 mmol) in dry DMF (1 mL) was added EDC.HCl (18 mg, 94 mol). The mixture was stirred overnight at rt and directly purified prep HPLC to give the bis TFA salt of the title compound (16 mg, 29%) as a yellow solid.

$^1H$ NMR (300MHz, DMSO-$d_6$) Shift = 7.96 (t, J=5.5 Hz, 1H), 7.59 - 7.51 (m, 3H), 7.49 - 7.43 (m, 2H), 7.34 - 7.28 (m, 2H), 7.23 - 7.10 (m, 4H), 7.07 - 6.98 (m, 2H), 6.78 - 6.68 (m, 1H), 5.03 (dd, J=5.5, 13.0 Hz, 1H), 3.87 (s, 2H), 3.61 - 3.22 (m, 14H), 2.96 - 2.34 (m, 9H), 2.07 - 1.93 (m, 1H), 1.89 - 1.69 (m, 2H).

$^{19}F$ NMR (282MHz, DMSO-$d_6$) Shift = -74.84, -117.44.

LC-MS $t_R$ = 5.30 min, m/z 935.

HRMS Calc'd for $C_{48}H_{49}ClFN_8O_9$: 935.328958; found: 935.329929.

**2-[2-(4-{2-[1-(5-chloro-1H-1,3-benzodiazol-2-yl)-5-hydroxy-3-[4-(trifluoromethyl)phenyl]-1H-pyrazol-4-yl]ethyl}phenoxy)ethoxy]-N-(2-{[2-(2,6-dioxopiperidin-3-yl)-1,3-dioxo-2,3-dihydro-1H-isoindol-4-yl](methyl)amino}ethyl)acetamide (FC-13887)**

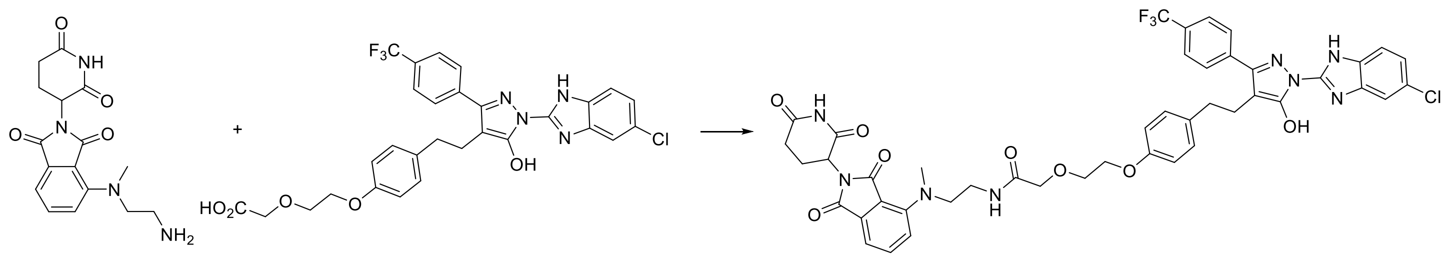

To a stirred solution of the HCl salt of 2-[2-(4-{2-[1-(5-chloro-1H-1,3-benzodiazol-2-yl)-5-hydroxy-3-[4-(trifluoromethyl)phenyl]-1H-pyrazol-4-yl]ethyl}phenoxy)ethoxy]acetic acid (18 mg, 28 µmol), the HCl salt of 4-[(2-aminoethyl)(methyl)amino]-2-(2,6-dioxopiperidin-3-yl)-2,3-dihydro-1H-isoindole-1,3-dione (11 mg, 30 µmol), HOBt.$H_2O$ (5 mg, 32 µmol) and i-$Pr_2NEt$ (30 µL, 0.17 mmol) in dry DMF (1 mL) was added EDC.HCl (11 mg, 57 mol). The mixture was stirred at rt for 1 d and purified by prep HPLC to give the bis TFA salt of the title compound (16 mg, 49%) as a yellow solid.

$^{1}H$ NMR (300MHz, DMSO-$d_6$) Shift = 7.91-7.79 (m, 4H), 7.72-7.64 (m, 1H), 7.61 - 7.52 (m, 2H), 7.27 - 7.21 (m, 2H), 7.20 - 7.15 (m, 1H), 7.06 - 6.99 (m, 2H), 6.77-6.70 (m, 2H), 5.13 - 5.00 (m, 1H), 3.95 (br. s., 2H), 3.73 (s, 2H), 3.63 - 3.47 (m, 4H), 3.43 - 3.25 (m, 2H), 2.99 (s, 3H), 2.97-2.65 (m, 5H), 2.58-2.38 (m, 2H), 2.01 - 1.80 (m, 1H).

$^{19}F$ NMR (282MHz, DMSO-$d_6$) Shift = -61.09, -74.88.

LC-MS $t_R$ = 5.23 min, m/z 913.

HRMS Calc'd for $C_{45}H_{41}ClF_3N_8O_8$: 913.268249; found: 913.268091.

**2-{2-[2-(3-{4-[1-(5-chloro-1H-1,3-benzodiazol-2-yl)-4-[2-(4-fluorophenyl)ethyl]-5-hydroxy-1H-pyrazol-3-yl]phenyl}propoxy)ethoxy]ethoxy}-N-{2-[2-(2-{[2-(2,6-dioxopiperidin-3-yl)-1,3-dioxo-2,3-dihydro-1H-isoindol-4-yl]amino}ethoxy)ethoxy]ethyl}acetamide (FC-13935)**

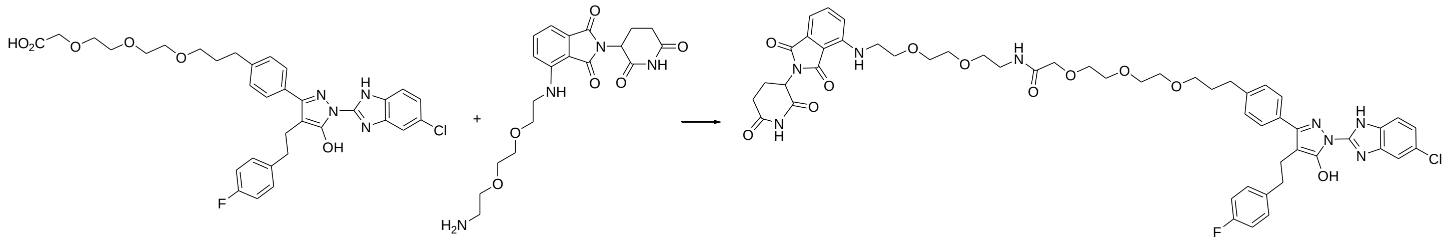

To a stirred solution of the HCl salt of 2-{2-[2-(3-{4-[1-(5-chloro-1H-1,3-benzodiazol-2-yl)-4-[2-(4-fluorophenyl)ethyl]-5-hydroxy-1H-pyrazol-3-yl]phenyl}propoxy)ethoxy]ethoxy}acetic acid (23 mg, 36 µmol), the HCl salt of 4-({2-[2-(2-aminoethoxy)ethoxy]ethyl}amino)-2-(2,6-dioxopiperidin-3-yl)-2,3-dihydro-1H-isoindole-1,3-dione (16 mg, 36 µmol), HOBt.$H_2O$ (6 mg, 40 µmol) and i-$Pr_2NEt$ (35 µL, 0.19 mmol) in dry DMF (1 mL) was stirred at rt for 1 d. The mixture was purified by prep HPLC to give the bis TFA salt of the title compound (7 mg, 15%) as a yellow solid.

$^{1}H$ NMR (300MHz, DMSO-$d_6$) Shift = 7.67 - 7.60 (m, 1H), 7.59 - 7.50 (m, 3H), 7.50 - 7.43 (m, 1H), 7.35 - 7.28 (m, 1H), 7.24 - 6.97 (m, 7H), 6.65 - 6.53 (m, 1H), 5.13 - 4.95 (m, 1H), 3.86 (s, 2H), 3.60-3.30 (m, 20H), 3.30 - 3.15 (m, 2H), 2.95 - 2.35 (m, 9H), 2.09 - 1.91 (m, 1H), 1.89 - 1.69 (m, 2H).

$^{19}F$ NMR (282MHz, DMSO-$d_6$) Shift = -74.80, -117.40.

LC-MS $t_R$ = 5.33 min, m/z 1023.

HRMS Calc'd for $C_{52}H_{57}ClFN_8O_{11}$: 1023.381387; found: 1023.381838.

**5-(4-{[1-(2-{2-[2-(3-{4-[1-(5-chloro-1H-1,3-benzodiazol-2-yl)-4-[2-(4-fluorophenyl)ethyl]-5-hydroxy-1H-pyrazol-3-yl]phenyl}propoxy)ethoxy]ethoxy}acetyl)piperidin-4-yl]methyl}piperazin-1-yl)-2-(2,6-dioxopiperidin-3-yl)-2,3-dihydro-1H-isoindole-1,3-dione (FC-14228)**

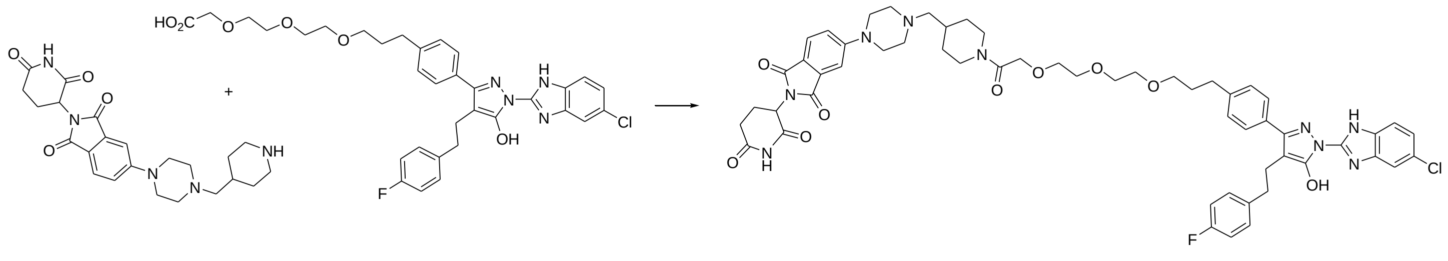

To a stirred solution of 2-{2-[2-(3-{4-[1-(5-chloro-1H-1,3-benzodiazol-2-yl)-4-[2-(4-fluorophenyl)ethyl]-5-hydroxy-1H-pyrazol-3-yl]phenyl}propoxy)ethoxy]ethoxy}acetic acid (12 mg, 18 µmol), the bis HCl salt of 2-(2,6-dioxopiperidin-3-yl)-5-{4-[(piperidin-4-yl)methyl]piperazin-1-yl}-2,3-dihydro-1H-isoindole-1,3-dione (11.5 mg, 22 µmol), HOBt.$H_2O$ (3 mg, 20 µmol) and i-$Pr_2NEt$ (25 µL, 0.14 mmol) in dry DMF (1 mL) was added EDC.HCl (7 mg, 37 mol). The mixture was stirred at rt for 1 d and purified by prep HPLC to give the tris HCl salt of the title compound (3 mg, 12%) as a solid.

$^1H$ NMR (300MHz, DMSO-$d_6$) Shift = 7.75 (d, J=8.3 Hz, 1H), 7.59 - 7.51 (m, 2H), 7.50 - 7.44 (m, 3H), 7.36 - 7.30 (m, 3H), 7.23 - 7.12 (m, 3H), 7.08 - 6.98 (m, 2H), 5.08 (dd, J=5.3, 12.7 Hz, 1H), 4.41 - 4.05 (m, 10H), 3.85-3.20 (s, 16H), 3.17 - 2.34 (m, 10H), 2.16 - 1.95 (m, 2H), 1.93 - 1.62 (m, 3H), 1.28 - 0.89 (m, 2H).

LC-MS $t_R$ = 4.52 min, m/z 1058.

HRMS Calc'd for $C_{56}H_{62}ClFN_9O_9$: 1058.433757; found: 1058.433249.

**3-[4-(2-{4-[1-(5-chloro-1H-1,3-benzodiazol-2-yl)-4-[2-(4-fluorophenyl)ethyl]-5-hydroxy-1H-pyrazol-3-yl]piperidin-1-yl}-2-oxoethoxy)-1-oxo-2,3-dihydro-1H-isoindol-2-yl]piperidine-2,6-dione (FC-14367)**

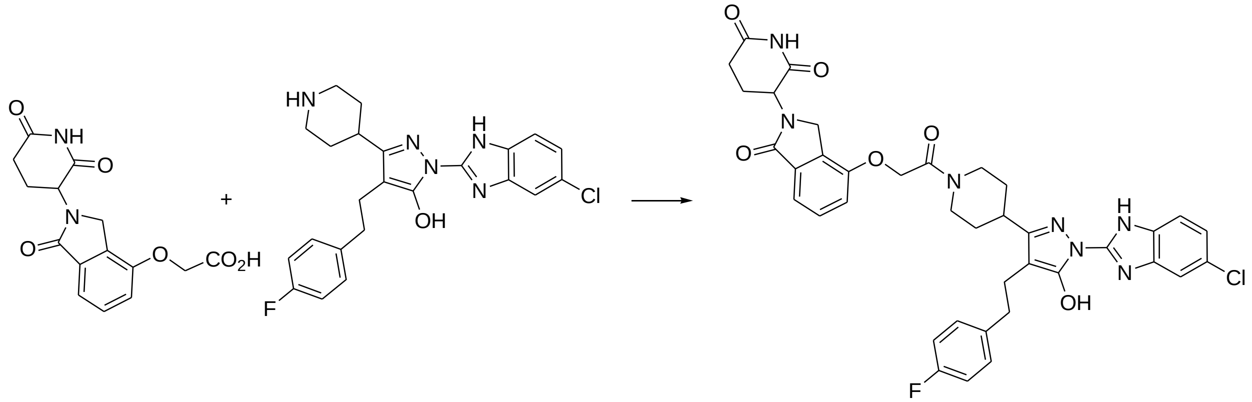

To a stirred solution of the HCl salt of 1-(5-chloro-1H-1,3-benzodiazol-2-yl)-4-[2-(4-fluorophenyl)ethyl]-3-(piperidin-4-yl)-1H-pyrazol-5-ol (29 mg, 57 µmol), 2-{[2-(2,6-dioxopiperidin-3-yl)-1-oxo-2,3-dihydro-1H-isoindol-4-yl]oxy}acetic acid (19 mg, 60 µmol), HOBt.$H_2O$ (9 mg, 60 µmol) and i-$Pr_2NEt$ (60 µL, 0.34 mmol) in dry DMF (1.5 mL) was added EDC.HCl (22 mg, 0.11 mmol). The mixture was stirred at rt for 18 h and purified by prep HPLC to give the TFA salt of the title compound (12 mg, 25%) as a white solid.

$^1H$ NMR (300MHz, DMSO-$d_6$) Shift = 10.99 (s, 1H), 7.56 - 7.46 (m, 2H), 7.46 - 7.41 (m, 1H), 7.34 - 7.29 (m, 1H), 7.26 - 7.02 (m, 6H), 5.19 - 4.89 (m, 3H), 4.49 - 4.15 (m, 4H), 3.97 - 3.77 (m, 1H), 3.14 - 2.97 (m, 1H), 2.78 (d, J=7.0 Hz, 3H), 2.66 - 2.33 (m, 4H), 2.08 - 1.90 (m, 1H), 1.88 - 1.55 (m, 2H), 1.53 - 1.32 (m, 2H).

$^{19}F$ NMR (282MHz, DMSO-$d_6$) Shift = -74.83, -117.46.

LC-MS $t_R$ = 4.63 min, m/z 740.

HRMS Calc'd for $C_{38}H_{36}ClFN_7O_6$: 740.239414; found: 740.239216.

**2-(4-{2-[1-(5-chloro-1H-1,3-benzodiazol-2-yl)-5-hydroxy-3-[4-(trifluoromethyl)phenyl]-1H-pyrazol-4-yl]ethyl}phenoxy)-N-(2-{[2-(2,6-dioxopiperidin-3-yl)-1,3-dioxo-2,3-dihydro-1H-isoindol-4-yl]amino}ethyl)acetamide (FC-14369)**

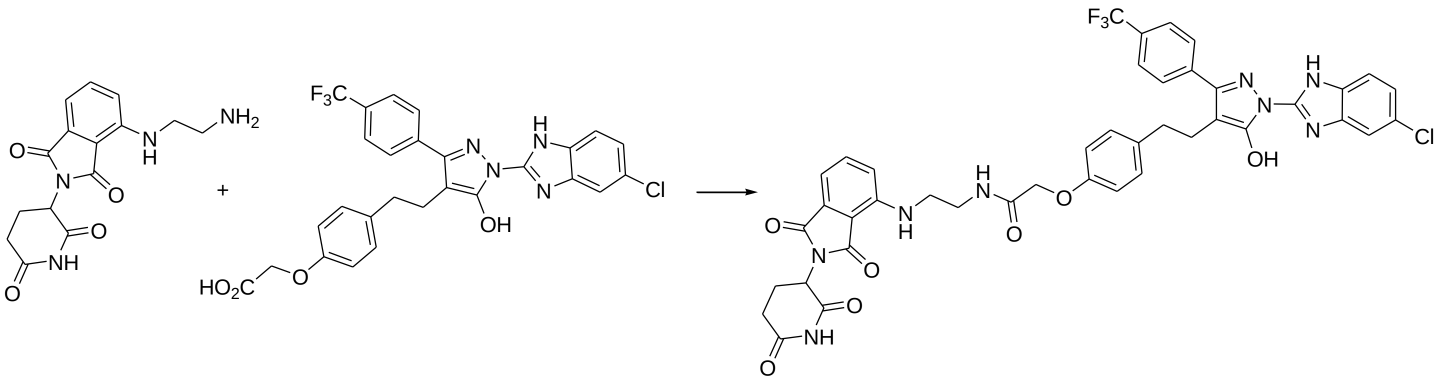

To a stirred solution of HCl salt 2-(4-{2-[1-(5-chloro-1H-1,3-benzodiazol-2-yl)-5-hydroxy-3-[4-(trifluoromethyl)phenyl]-1H-pyrazol-4-yl]ethyl}phenoxy)acetic acid (33 mg, 56 µmol), the HCl salt of 4-[(2-aminoethyl)amino]-2-(2,6-dioxopiperidin-3-yl)-2,3-dihydro-1H-isoindole-1,3-dione (21 mg, 60 µmol), HOBt.$H_2O$ (8 mg, 53 µmol) and i-$Pr_2NEt$ (60 µL, 0.34 mmol) in dry

DMF (1.5 mL) was added EDC.HCl (22 mg, 0.11 mmol). The mixture was stirred at rt for 18 h and directly purified by prep HPLC to give the TFA salt of the title compound (29 mg, 54%) as a yellow solid.

$^{1}$H NMR (300MHz, DMSO-$d_6$) Shift = 8.35 - 8.21 (m, 1H), 7.89 - 7.81 (m, 2H), 7.80-7.77 (m, 2H), 7.61 - 7.50 (m, 2H), 7.28 - 7.21 (m, 1H), 7.20 - 7.15 (m, 1H), 7.08 - 6.96 (m, 2H), 6.84 - 6.69 (m, 3H), 5.04 (dd, J=5.5, 12.5 Hz, 1H), 4.39 (s, 2H), 3.47 - 3.22 (m, 4H), 2.99 - 2.78 (m, 1H), 2.73 (s, 4H), 2.61 - 2.35 (m, 2H), 2.09 - 1.92 (m, 1H).

$^{19}$F NMR (282MHz, DMSO-$d_6$) Shift = -61.09, -74.78.

LC-MS $t_R$ = 5.28 min, m/z 855.

HRMS Calc'd for $C_{42}H_{35}ClF_3N_8O_7$: 855.226384; found: 855.224689.
